# EAGLE: an algorithm that utilizes a small number of genomic features to predict tissue/cell type-specific enhancer-gene interactions

**DOI:** 10.1101/781427

**Authors:** Tianshun Gao, Jiang Qian

## Abstract

Long-range regulation by distal enhancers is crucial for many biological processes. The existing methods for enhancer-target gene prediction often require many genomic features. This makes them difficult to be applied to many cell types, in which the relevant datasets are not always available. Here, we design a tool EAGLE, an <u>e</u>nhancer <u>a</u>nd <u>g</u>ene <u>l</u>earning <u>e</u>nsemble method for identification of Enhancer-Gene (EG) interactions. Unlike existing tools, EAGLE used only six features derived from the genomic features of enhancers and gene expression datasets. Cross-validation revealed that EAGLE outperformed other existing methods. Enrichment analyses on special transcriptional factors, epigenetic modifications, and eQTLs demonstrated that EAGLE could distinguish the interacting pairs from non- interacting ones. Finally, EAGLE was applied to mouse and human genomes and identified 7,680,203 and 7,437,255 EG interactions involving 31,375 and 43,724 genes, 138,547 and 177,062 enhancers across 89 and 110 tissue/cell types in mouse and human, respectively. The obtained interactions are accessible through an interactive database enhanceratlas.org. The EAGLE method is available at https://github.com/EvansGao/EAGLE and the predicted datasets are available in http://www.enhanceratlas.org/.

**Author summary:** Enhancers are DNA sequences that interact with promoters and activate target genes. Since enhancers often located far from the target genes and the nearest genes are not always the targets of the enhancers, the prediction of enhancer-target gene relationships is a big challenge. Although a few computational tools are designed for the prediction of enhancer-target genes, it’s difficult to apply them in most tissue/cell types due to a lack of enough genomic datasets. Here we proposed a new method, EAGLE, which utilizes a small number of genomic features to predict tissue/cell type-specific enhancer-gene interactions. Comparing with other existing tools, EAGLE displayed a better performance in the 10-fold cross-validation and cross-sample test. Moreover, the predictions by EAGLE were validated by other independent evidence such as the enrichment of relevant transcriptional factors, epigenetic modifications, and eQTLs.

Finally, we integrated the enhancer-target relationships obtained from human and mouse genomes into an interactive database EnhancerAtlas, http://www.enhanceratlas.org/.

## Introduction

Enhancers function as distal cis-regulatory elements for the regulation of target gene expression (1). They are tissue/cell type-specific and usually display in clusters of redundant elements to regulate the gene expression (2). Many approaches were developed to infer enhancer activity on a genome-wide scale. For example, mapping the genome-wide locations of P300, an enzyme that is a good indicator of enhancers, can help to identify enhancers in a particular cell type (3). Specific histone modifications (e.g. H3K27ac and H3K4me1) were used to predict enhancer activity (4). Chromatin accessibility measured by DNase I hypersensitivity or ATAC-seq is also a good measurement of active enhancers because enhancers are located in open chromatin regions (5, 6). Transcribed enhancer sequences (eRNAs) were also used to measure the enhancer activity in different cell types (1). A variety of these datasets were generated in many cell types (7).

Identification of enhancer-target interactions is much more challenging than the measurement of enhancer activity (8–11). Since active enhancers interact with promoters in 3D space, the interactions detected by Hi-C or ChIA-PET were often used to predict enhancer-target relationships (12, 13). However, Hi-C and ChIA-PET are still difficult and expensive assays to perform in the laboratories for detection of enhancer-promoter loops in most tissue/cell types. Furthermore, the resolution of the interactions detected using Hi-C was often low. The locations of enhancers from the target genes were often in the range of 2kb to 10kb (10). Therefore, in silico prediction based on the ChIA-PET or Hi-C training model would be an economic method to identify tissue/cell-specific enhancer-target interactions in many tissue/cell types.

Several computational approaches have been developed to predict enhancer-target relationships (3, 8–11, 14, 15). Assigning the nearest gene to enhancer is the simplest approach to predict these relationships. However, the approach ignored the long-range interactions between enhancers and promoters (16). The 5C experiment demonstrated that only 7% of the enhancer-promoter interactions owned the nearest genes (16). Correlated activities between enhancers and promoters based on chromatin activity or histone modifications were also used to determine the relationships (14, 15). Deep analyzing published Hi-C data (e.g. PSYCHIC) was another method to recognize high-quality enhancer-promoter interactions (17). Recently, several machine-learning approaches (e.g. IM-PET, Ripple, TargetFinder, and JEME) were developed, which integrated multiple genomic features to predict the relationships (8–11). However, these approaches were only applied in human and trained with many extra features such as histone modification, transcription factor (TF) binding, evolution, and chromatin accessibility. Therefore, it is hard to apply these methods to other species or tissue/cell types with few available features.

In this work, we develop a method called EAGLE, an enhancer and gene learning ensemble method, to predict the enhancer-target relationships. Our approach utilizes only six genomic features so that we can apply the method to many cell types. Some of the features were never used in previous algorithms. For example, we calculated the correlation between the pairwise enhancer activities across cell types based on the observation that many enhancers cooperate to co-regulate target genes. We also considered the numbers of enhancers and genes between the enhancer of interest and the target gene. These features improve the performance of our prediction. Finally, we applied EAGLE to 110 and 89 cell types in human and mouse, respectively. The predicted relationships are integrated into EnhancerAtlas.org so that users can retrieve and visualize them in the enhancer browser.

## Results

### A new approach to predict enhancer-target relationships

To predict interacting enhancer-target pair in a particular cell type, we built a machine learning approach called EAGLE using the 6 features described below (Fig 1). The input of the model is the enhancer annotation and gene expression data. The enhancer annotation was obtained from our previous work, in which we integrated multiple genomic datasets to derive a set of reliable enhancer annotation in different tissue/cell types (18). The training data were defined by ChIA-PET datasets in human or Hi-C datasets in the mouse. We defined the positives as the pairs overlapping with ChIA-PET or Hi-C interactions, while the negatives were set as the pairs that had no overlaps with them. Note that for both positive and negative pairs, the distances of their EG were limited within 1Mbp and the selected enhancers and promoters were both active. We tried different machine learning methods (e.g. linear regression, SVM, KNN, Discriminant, Decision tree, and boosting trees) and chose the learning ensemble boosting method, which was with the highest performance among all (Fig S1).

**Fig 1.**
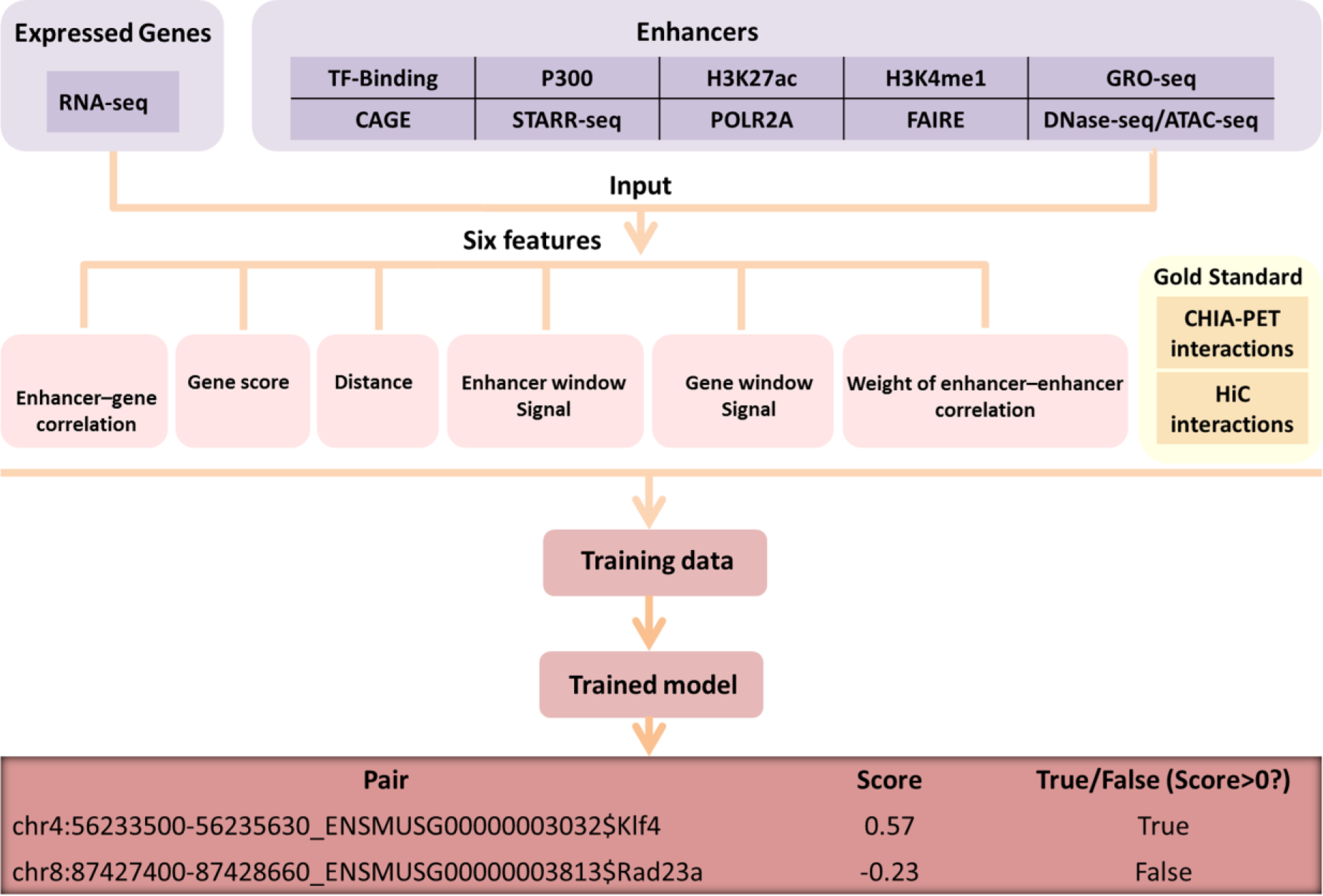
Overview of the EAGLE pipeline. Enhancers were obtained by integrating diverse high-throughput datasets and the expressed levels were estimated using RNA-seq data. We utilized six features based on the information of enhancers and gene expression. ChIA-PET or Hi-C datasets were used to define positive and negative EG pairs. Using the labeled pairs, we trained an ensemble classifier, EAGLE, which could predict enhancer-target interactions measured by prediction probabilities.

### Six genomic features used to predict EG interactions

To make the method applicable to as many tissue/cell types as possible, we did not use the auxiliary information (e.g. histone modification, TF bindings). Only six features were used in EAGLE. These features were tested on 71,118 positive, 71,118 random and 71,118 negative EG pairs defined by ChIA-PET data in K562 respectively, and 9732 positive, 9,732 random and 9,732 negative ones in GM12878 respectively (Fig 2 and Fig S2). Here the positive EG pairs were defined as the EG interaction candidates that overlapped with ChIA-PET interactions, while the negative EG pairs were the EG interaction candidates that have no any overlaps with ChIA-PET. To compare with positives and negatives, a certain number of randomly selected EG interaction candidates were taken as the “random” group.

**Fig 2.**
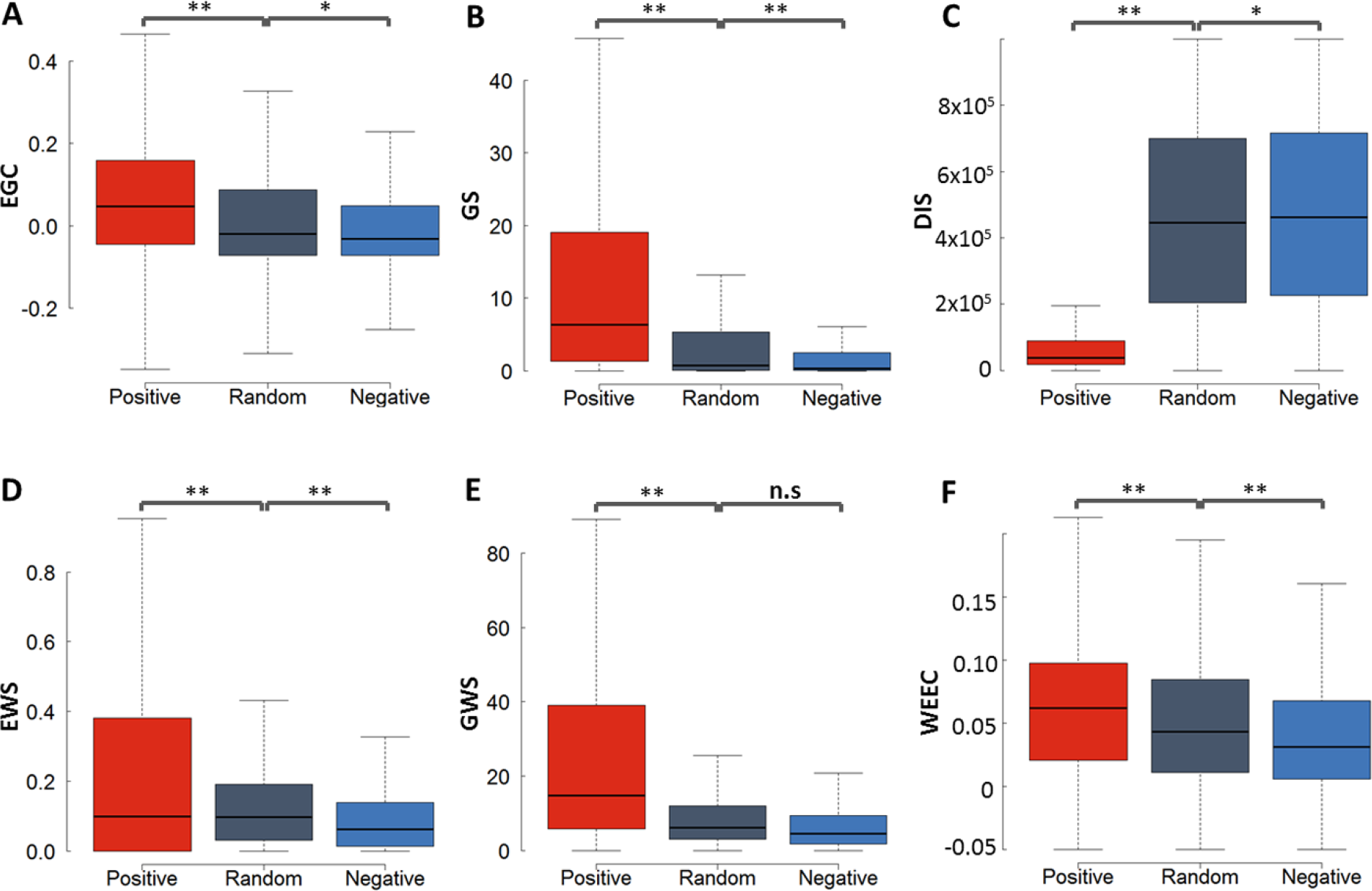
Six discriminative features based on the ChIA-PET data from K562. (A) Enhancer activity and gene expression profile correlation (EGC). (B) Gene score (GS) from the RNA-seq data. (C) Distance (DIS) between enhancer and gene in a pair. (D) Enhancer window signal (EWS) measuring the mean enhancer signal in the region between enhancer and promoter (E) Gene window signal (GWS) evaluating the mean gene expression level in the region between enhancer and promoter (F) The weight of enhancer-enhancer correlation (WEEC). The positive, negative and random EG pairs were obtained from ChIA-PET dataset in K562. The *P* values were calculated using the Student *t*-test. \**P* < 0.01; \*\**P* < 1e-16; n.s. not significant.

#### Enhancer activity and gene expression profile correlation (EGC)

We expect that the activity of an enhancer and the expression level of the target gene have a certain degree of correlation with each other. We used the score for the enhancer annotation as a proxy of enhancer activity. The expression levels of genes were based on RNA-seq measurement. The correlation was calculated across 110 and 89 cell types for human and mouse, respectively. As shown in Figure 2A, the correlations for the interacting (positive) EG pairs were significantly higher than those for non-interacting (negative) EG pairs. The random EG pairs, which included both positive and negative pairs, have intermediate correlation level.

#### Gene score (GS)

We expect that a real EG interaction indicates a strong activity in the interacting gene. Therefore, the gene score, which reflected the expression level of the target genes in a particular cell type, is also a useful feature to determine the active enhancer-target gene relationship. As expected (Fig 2B), the genes interacting with enhancers have a higher expression level than those without interaction with enhancers.

#### Distance (DIS)

The linear genomic distance played an important role in defining the enhancer-target pairs. Generally, positive EG pairs have much shorter distances than negative EG pairs (Fig 2C). In K562 and MCF-7, the median genomic distances of the positive pairs are 46,934 bp in K562 and 37,556 bp in MCF-7, while the median distances of the negatives are 468,448 bp in K562 and 490,667 bp in MCF-7 (Fig 2C and Fig S2). The distribution for EG distances in positives is also different from the one in negatives (Fig S3A). However, the distances in real EG pairs are still much larger than those in pairs with the nearest genes (*p*=0, t-test), suggesting that we cannot predict the EG pairs simply based on the nearest genes (Fig S3E). These results indicated that the genomic distance between the interacting enhancer and promoter was a very discriminative feature that can distinguish the positives from negatives or pairs with the nearest genes.

#### Enhancer window signal (EWS)

We defined the EWS as the mean enhancer signal in the window between EG. Since most enhancers do not interact with the nearest genes, we wonder whether the information of other enhancers located between the enhancer and gene of one EG pair plays an important role. We observed that the signal of enhancers between EG increases with the enlargement of the EG distance, so we normalized the enhancer signal between EG by the distance. Interestingly, the normalized EWS in true EG pairs (positives) is significantly higher than that in false EG pairs (negatives) (Fig 2D).

#### Gene window signal (GWS)

Similarly, we also calculated the gene activity between the enhancer and gene of each pair. GWS is the summation of gene expression level divided by the distance between EG. We discovered that the GWS for interacting pairs is much higher than those in non-interacting pairs (Fig 2E).

#### Weight of enhancer-enhancer correlations (WEEC)

Multiple enhancers often cooperated to co-regulate the target genes (19). We calculated the correlation coefficient between the enhancers across all tissue/cell types. As shown in Figure 2F, interacting enhancers tend to have a higher correlation with other enhancers than the non-interacting enhancers.

### Performance evaluation of EAGLE

To evaluate the performance of EAGLE, we performed both self-testing and across-sample testing. The area under the receiver operating characteristic (AUROC) curve and the area under the Precision-Recall (AUPR) curve were both used to measure the performance (20). In the self-testing, we used one-half of the data for training and the other half for testing. This test was applied for K562 and MCF-7, respectively. The performances with the AUROC and AUPR values in K562 reached 95.65% and 95.28%, respectively (Fig 3). We also developed the model in MCF-7 and the performance could reach 93.25% and 91.98% for AUROC and AUPR, respectively (Fig 3). For the cross-sample validation, K562 data was trained to build the prediction model and GM12878 was used for testing. EAGLE also has an excellent performance with 93.38% for AUROC and 92.36% for AUPR in across-sample tests (Fig S4A). Similar results were obtained if we used other cell lines for training (Fig S5). EAGLE also works well with the unbalanced datasets. For example, using unbalanced data with a ratio of 1:5 between positives and negatives in GM12878, EAGLE still got good performances of 93.27% and 72.89% for AUROC and AUPR, respectively (Fig S4B).

**Fig 3.**
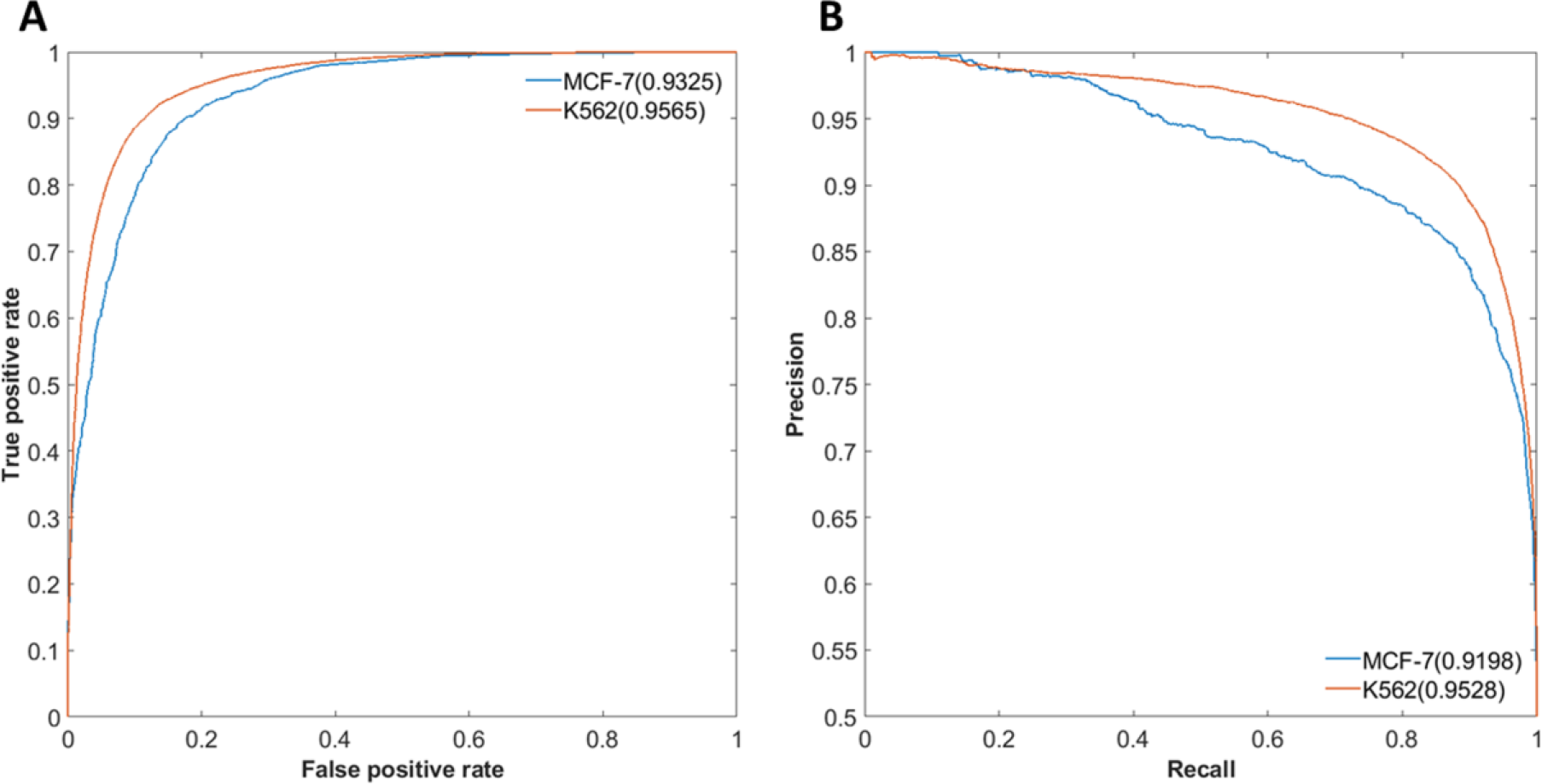
Performance evaluation of EAGLE. We trained an ensemble classifier to predict EG pairs. (A) Performance measurement of self-testing in K562 and MCF-7 by ROC curves. (B) Performance measurement of self-testing in K562 and MCF-7 by PR curves. In each cell line, one half of the data was taken for training, while the other half was used for testing. The performance was measured as the area under ROC or PR curves.

We then compared EAGLE with other four existing methods: JEME, IM-PET, TargetFinder, and Ripple. All the methods built the models using the ChIA-PET or Hi-C data from K562 (**See methods**). This comparison was made on the predictions in GM12878. EAGLE has the AUROC of 93.08%, while JEME, Ripple, IM-PET and TargetFinder have the corresponding values of 90.77%, 87.05%, 78.77%, and 83.39% respectively (Fig 4A). Similarly, EAGLE has a better performance in terms of the PR curve (Fig 4B). The results of other cell lines also demonstrated that EAGLE outperformed other existing methods (Fig S6).

**Fig 4.**
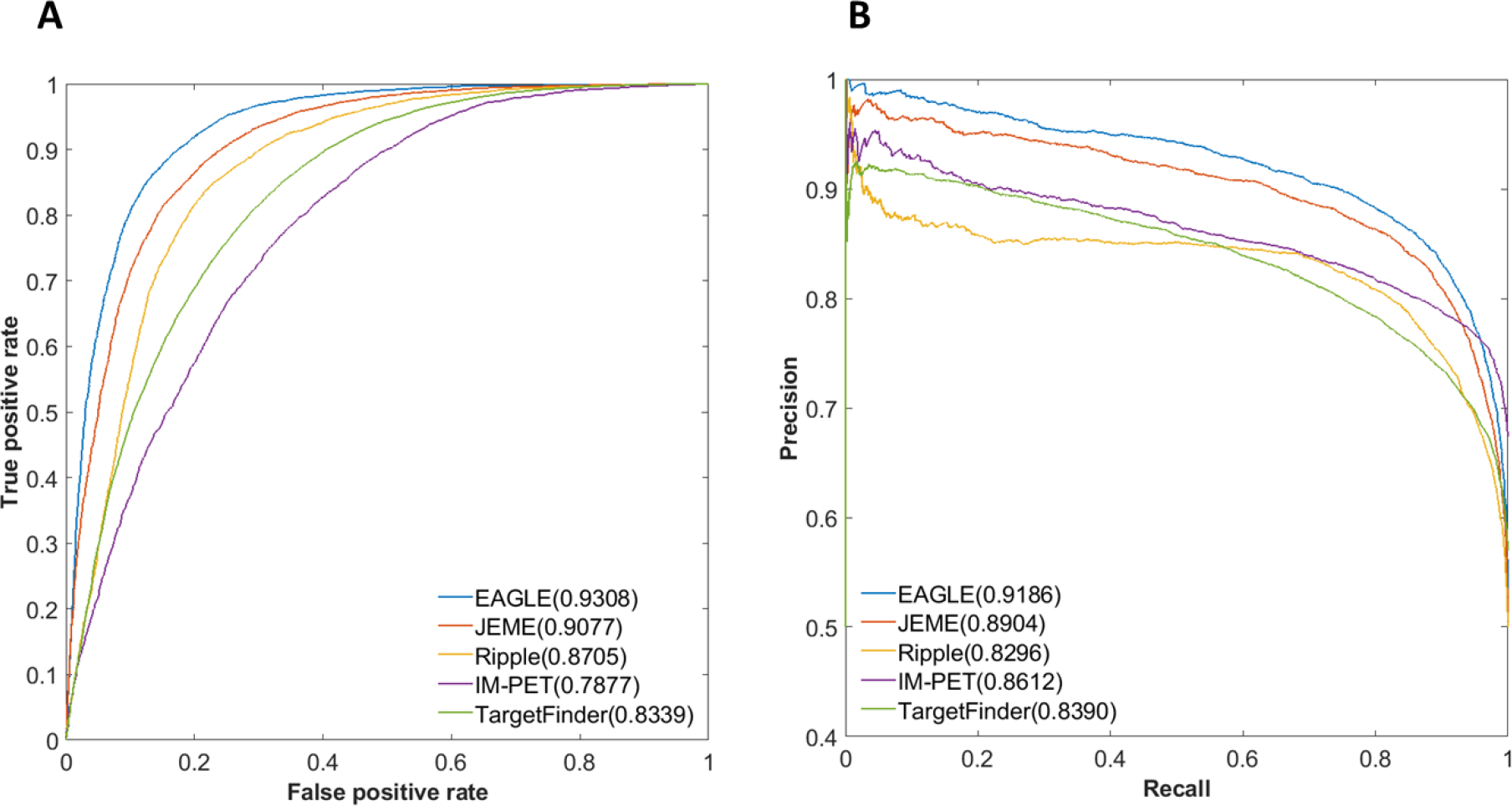
Performances of EG prediction tools based on cross-sample validation. (A) Relative AUROCs for all five methods (B) AUPRs for these methods. The model was trained based on K562 dataset while the prediction was made in GM12878 (see Methods).

### Importance of each feature in each cell type

It is clear that each feature contributes to the prediction. With an increasing number of features, the prediction performance increases (Fig S4 and S7). We then calculated the importance of each feature in four cell lines with enough ChIA-PET data for training. Using the permutation of out-of-bag predictor observation, which is similar to leave-one-out method, we estimated the relative importance of the features in each cell (Fig 5). Each feature was robust with effective importance (>=0.05) in each cell. Interestingly, some features performed very different importance in different cells. For example, WEEC has small importance with 0.05 in MCF-7, while its importance reached 0.18 in K562 (Fig 5).

**Fig 5.**
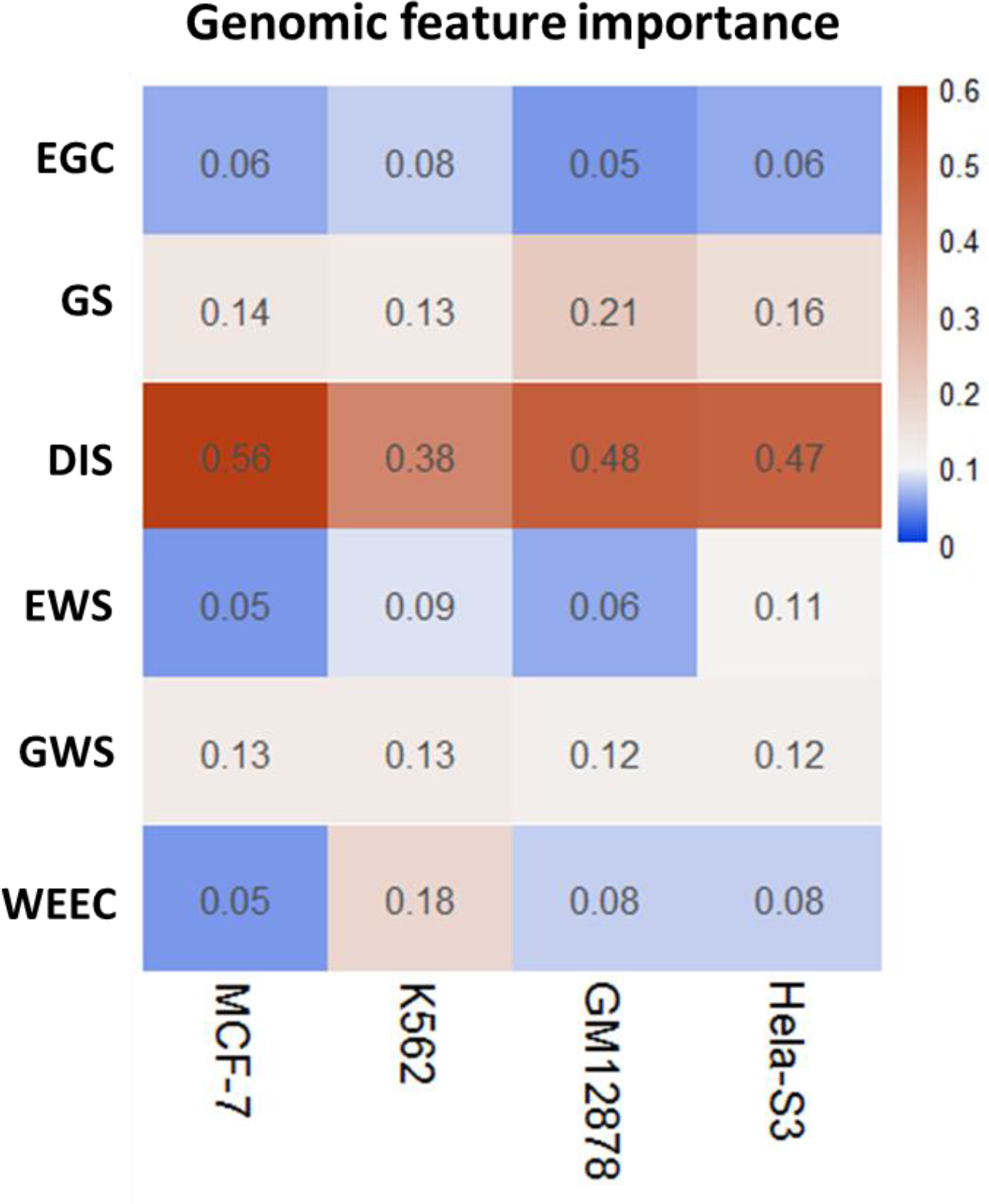
Predictive importance of training features across four common cell lines. The method permuting out-of-bag predictor observation was adopted to evaluate the relative importance of each feature. All features showed good robustness with effective importance (>=0.05) across all cell lines. Some features such as DIS contribute >=38% in all cells. However, some features such as WEEC and EWS have different importance in different cells.

### eQTL enrichment in putative EG interactions

We then used independent genomic information to validate the prediction. We used a series of genomic variables that were not used as features for prediction and compared these variables between the predicted positives and negatives. The first genomic variable we used is the eQTLs from GTEx portal (https://gtexportal.org) (21), which are the genetic variants that are likely to regulate the gene expression. Since eQTLs connected cis-regulatory elements to target genes, we asked whether the relationships identified by the eQTLs could be recovered by our prediction. We first collected eQTLs data from eight tissue types, including spleen, skeletal muscle, pancreas, ovary, lung, liver, left ventricle, and HMEC. These are the tissue types that overlap between our 70 predicted tissue/cells and 48 tissues with eQTL data. We calculated the percentage of predicted enhancer-target pairs that contain the eQTL relationships and compared the percentages between interacting and non-interacting enhancer-target pairs. We found that the predicted interacting pairs have a much higher percentage of containing eQTL relationships than the non-interacting pairs (Fig 6A). For example, in the spleen, 2.9% and 0.2% of enhancer-target pairs were reproduced by eQTLs in positive and negative datasets, respectively. Similarly, in the pancreas, 18.3% and 1.5% of enhancer-target pairs were reproduced by eQTLs in predicted positive and negative datasets, respectively. Furthermore, the eQTLs overlapping with non-interacting enhancer-target pairs are much less statistically significant than those overlapping with interacting enhancer-target pairs (Fig 6B). If we ignore the tissue specificity of the eQTLs and combined the eQTLs from all 48 tissues, we found that these combined eQTLs were enriched in interacting enhancer-target pairs for 70 human cell lines/tissues (Fig 6C).

**Fig 6.**
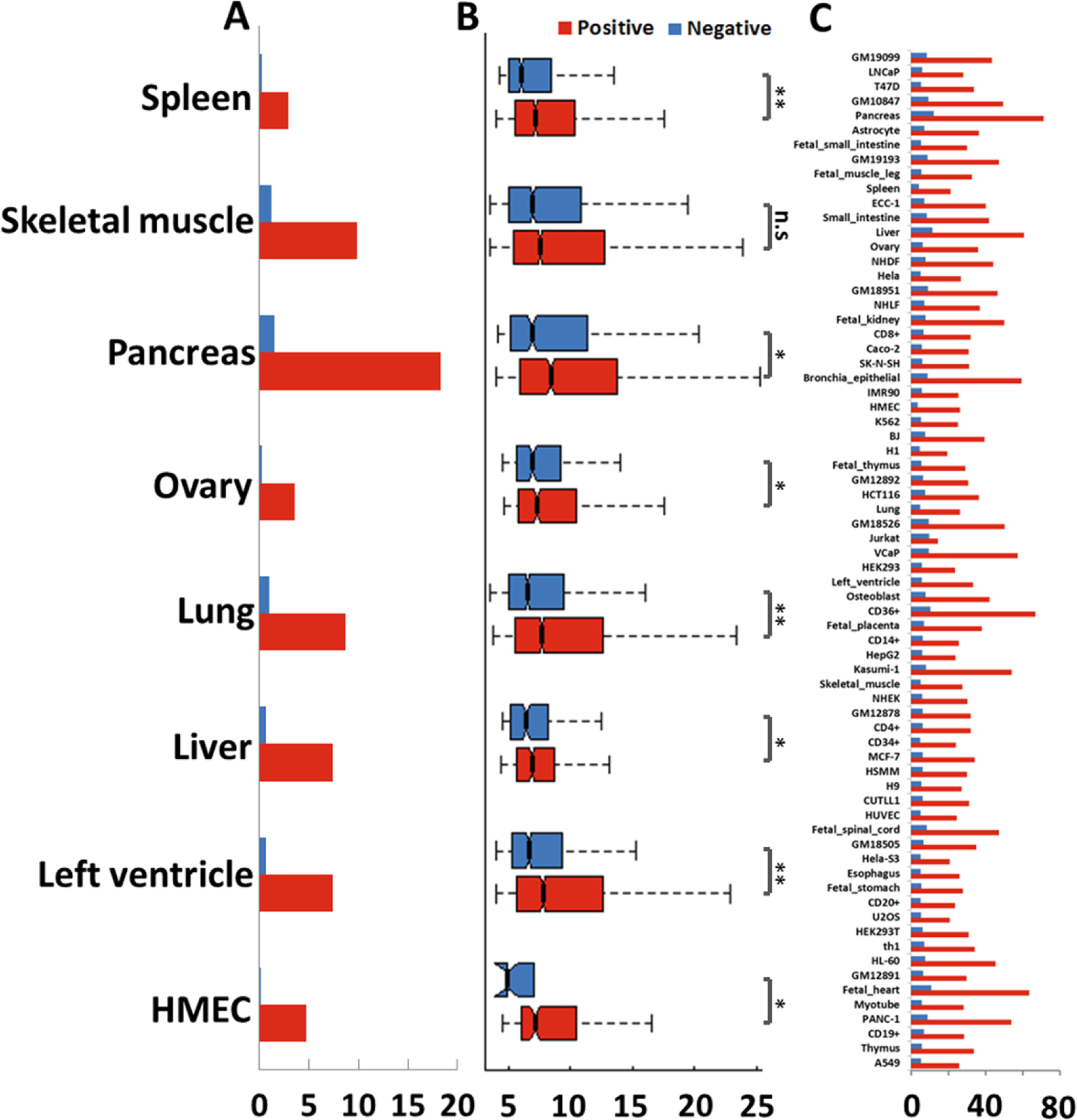
Predicted EG interactions are enriched for eQTLs. (A) Percentages of tissue-specific eQTLs overlapping with interacting and non-interacting pairs, respectively. The eQTL datasets were from the corresponding tissue types that we used to predict enhancer-target gene pairs. (B) Significance (p-values) of eQTLs overlapping with interacting and non-interacting enhancer-target pairs. \**P* < 0.01; \*\**P* < 1e-16; n.s. not significant. (C) Percentages of interacting and non-interacting pairs overlapping with eQTLs combined from 48 tissues.

### Validation of predicted interactions using genomic features

We then assessed whether the enhancers interacting with promoters and those not interacting with promoters have distinct characteristics. We examined a series of genomic features and compared their intensities (or frequencies) between interacting and non-interacting pairs in GM12878. For the enhancers, we examined three features H3K27ac, H3K4me1, and H3K9ac, while for the promoters we examined H3K4me3 (Fig 7). These histone marks represent active transcriptional activation (9, 10, 22), while CTCF and RAD21 are involved in the genomic looping that connects enhancers and their target genes (23). The intensities of the three histone marks on enhancers were higher in the interacting enhancers than those in the non-interacting enhancers. Similarly, CTCF, RAD21, and H3K4me3 occurred more often at interacting promoters than non-interacting promoters (Fig 7). Taken together, histone marks and relevant factors suggested that our prediction of enhancer-target relationships were likely biologically functional.

**Fig 7.**
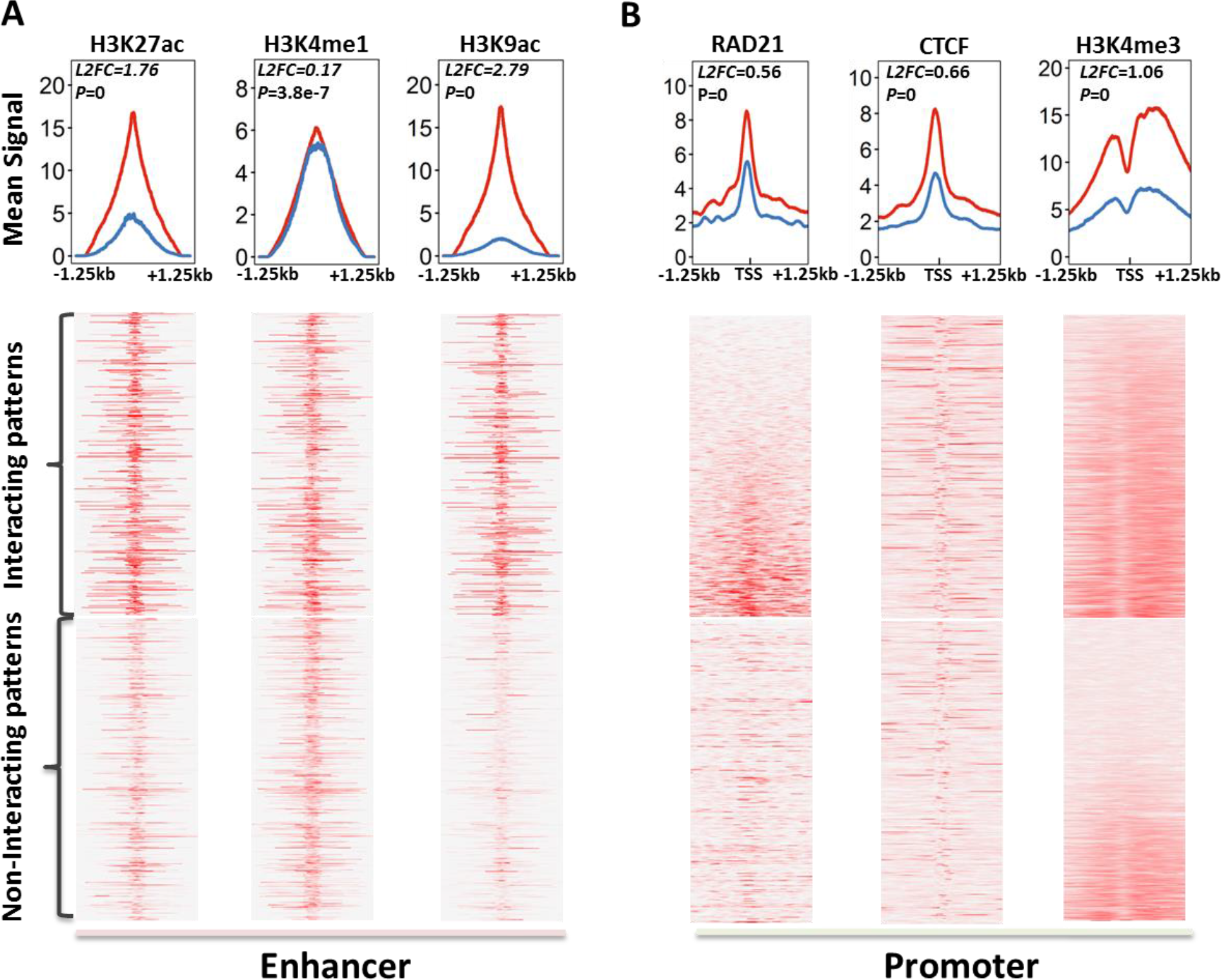
Chromatin states of interacting or non-interacting enhancers/promoters marked by related TFs or epigenetic modifications. (A) Density analysis and heatmaps for enhancers using enhancer marks (H3K4me1, H3K27ac, and H3K9ac). (B) Density analysis and heatmap for promoters using promoter marks (RAD21, CTCF, and H3K4me3). Total 7531 interacting and 1491 non-interacting enhancers in GM12878 were used for this analysis. Red lines marked the mean signal of the interacting enhancers (A) or promoters (B), while blue lines labeled the non-interacting elements. The p-value and log base 2 fold change (L2FC) in each plot indicated the statistically significant difference between interacting elements and non-interacting ones.

### Application to mouse and human tissue/cell types

We applied the EAGLE to predict enhancer-target relationships in mouse cells/tissues. For this purpose, we first determined the enhancer consensus in mouse cells/tissues by integrating various genomic datasets using the approach we developed (18) (Fig 8A). More than 6,000 high-throughput datasets were collected from ten high-through approaches (“Histone”, “TF-Binding”, “DHS”, “FAIRE”, “CAGE”, “EP300”, “POL2”, “MNase”, “GRO-seq”, and “STARR”) across 156 cell/tissue types (See Table S1 and data source in link http://www.enhanceratlas.org/download2.php). Many high-through approaches have been applied in many tissue/cells (Fig 8B). To ensure a high quality of enhancer annotation, only cell/tissue types with at least three independent experiments were selected for enhancer prediction (Fig 8C). By cross-validating the datasets and assessing the data quality for each cell/tissue types, we identified total 2,811,699 enhancers for 156 cell types.

**Fig 8.**
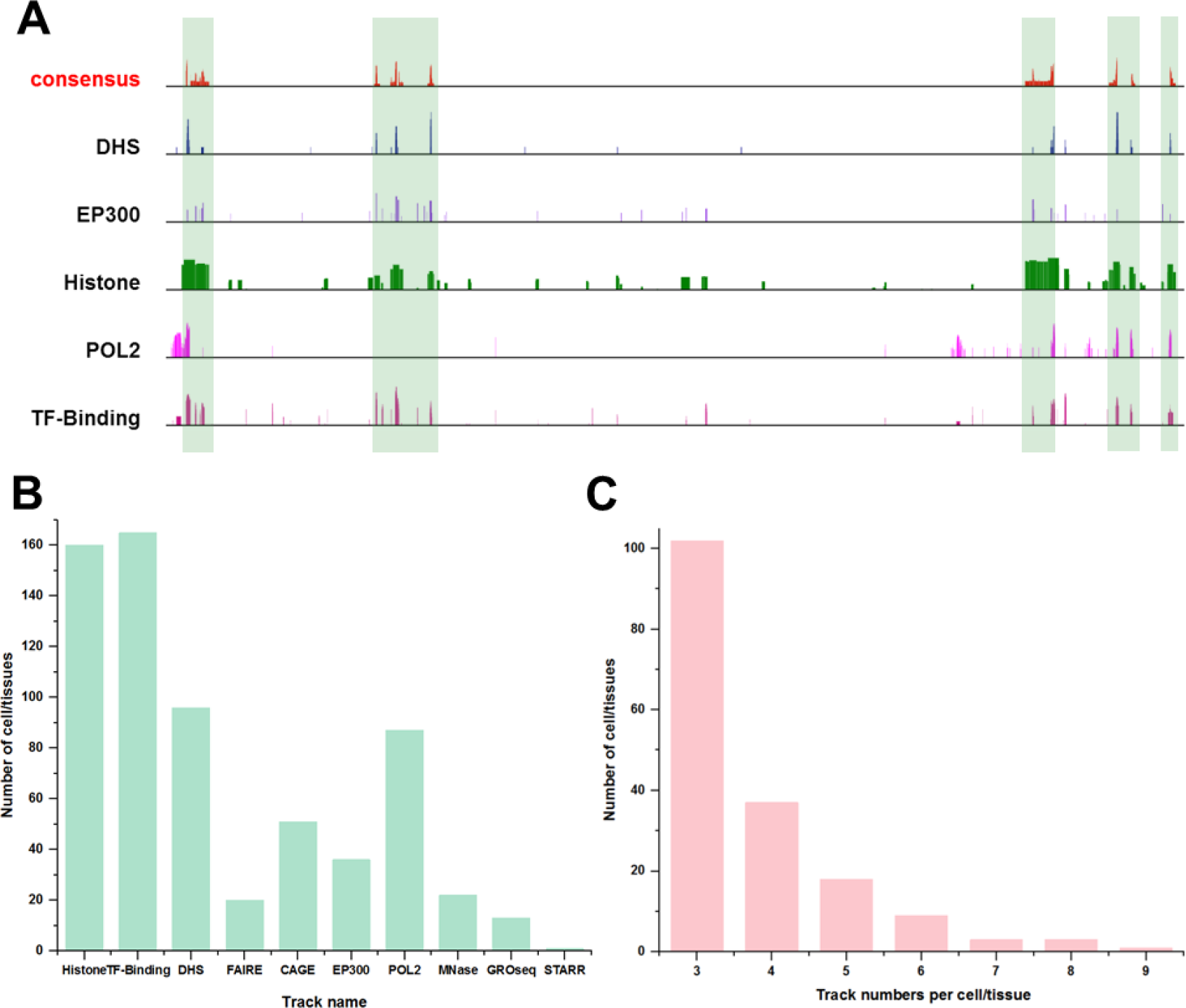
Enhancer consensus annotation in mouse cells. (**A**) Consistency and discrepancies in enhancer annotation. Vertical bars mark the enhancers supported by many tracks. Note that many regions are only supported by one or a few tracks. (**B**) Number of tissue/cell types that contain certain dataset types. Some technologies were more widely used than others for enhancer identification. (**C**) The number of cell/tissue types in function of the number of independent tracks. Many cell/tissue types include a few tracks (e.g. 3 or 4), while a few cell/tissue types have many tracks (e.g. 8 or 9).

Of the 156 cell types, 89 were found with the RNA-seq data available. We then predicted enhancer-target relationships in these cell types using EAGLE. We used the lung Hi-C data to train the model. The selected genomic features showed a significant difference between positives and negatives (Fig S8). In the self-testing, the performance of the model trained by mouse lung achieved 91.57% and 87.95% measured by AUROC and AUPR, respectively. The across-sample test on spleen also displayed high performances of AUROC and AUPR as 93.34% and 91.96%, respectively (Fig 9A, Fig S9). With this model, we predicted total 7,680,203 relationships involving 31,375 genes and 138,547 enhancers in the 89 cell types. On average, 86,294 relationships were identified in each cell type.

**Fig 9.**
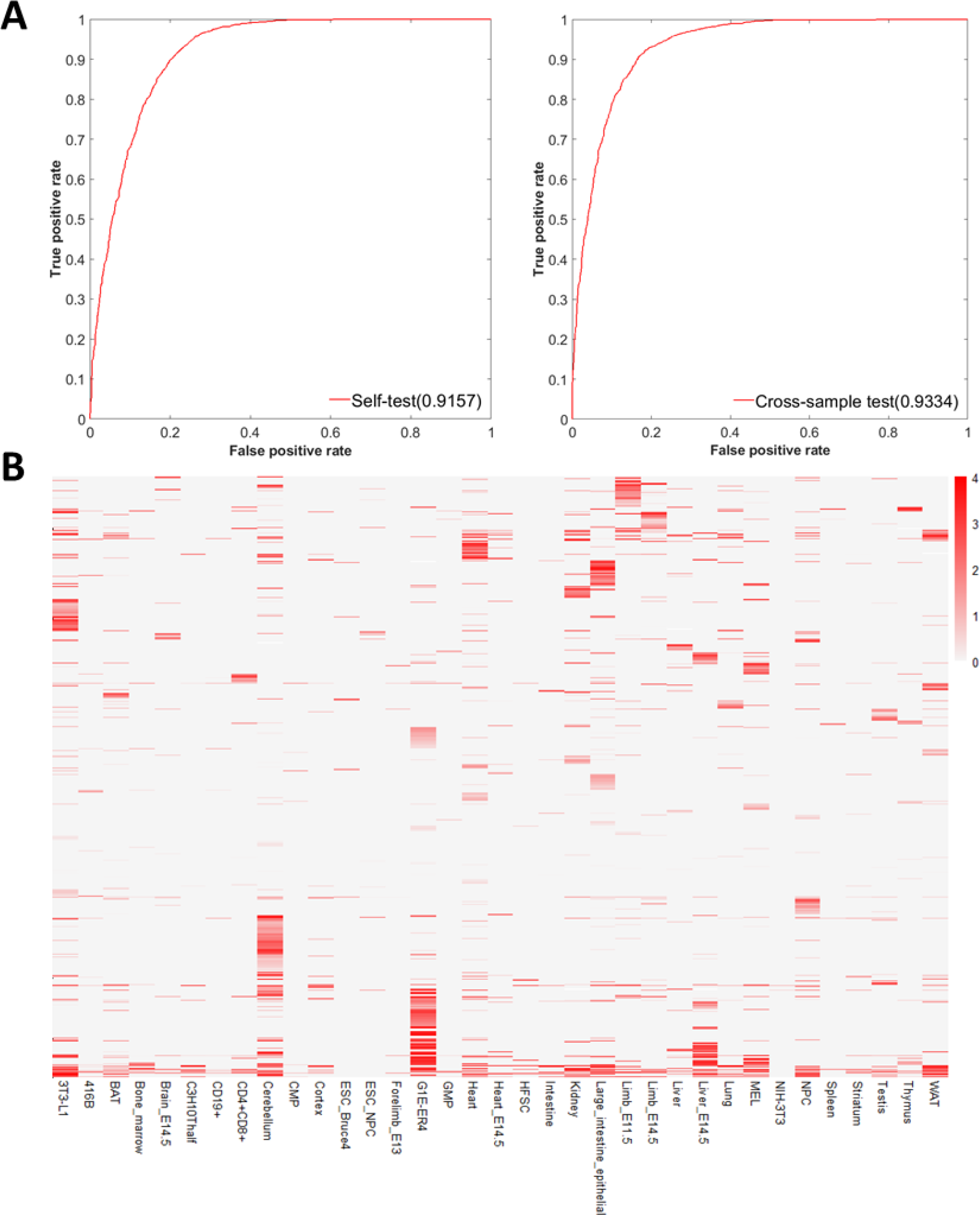
Application of EAGLE to mouse tissue/cells. (A) Performance of EAGLE model in the mouse by self-testing and cross-sample test. (B) Enhancer-target relationships across 35 representative cell types in the mouse. The EAGLE model in the mouse was trained in the lung. In the heatmap, each row represents an enhancer-target interaction, while each column is one particular cell type. The color represents the prediction confidence score for this interaction. On average, 95,723 relationships were identified in each cell type. Majority of the relationships were tissue-specific.

Similarly, we applied EAGLE to the human genome. We have identified 2,534,123 enhancers in 105 human cell types in our previous work (18). We then used EAGLE to predict enhancer-target relationships. In total, 7,437,255 enhancer-target gene relationships involving 43,724 genes and 177,062 enhancers were predicted in 110 tissue/cell types. These enhancer-target relationships can be queried and visualized in EnhancerAtlas.org.

We examined the enhancer-target relationships across 35 representative cell types in the mouse. As demonstrated in Figure 9B, the majority of the relationships were indeed tissue specific. Specifically, 53.0% and 19.2% of interactions occurred in one and two cell types, respectively.

We performed the similarity analysis and identified some patterns of interactions across cell-types and tissues (Fig. 10). We found that similar tissue types tend to have similar EG interactions. For example, the blood cell lines showed higher similarities among themselves, while other tissues showed lower similarities with the other tissue/cell lines.

**Fig 10.**
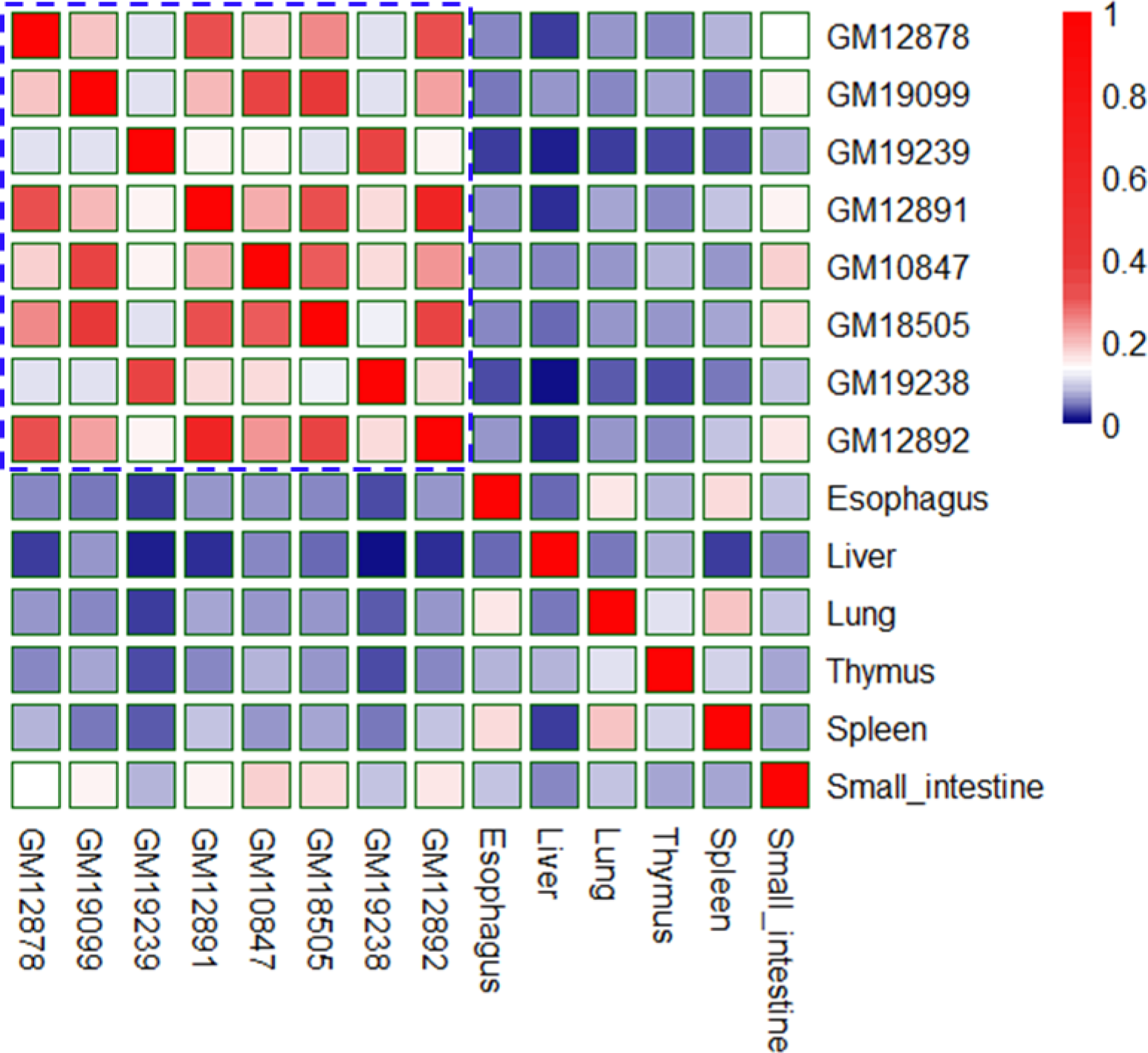
Patterns of predicted EG interactions across tissue/cell-types. Similarities were measured by Jaccard index across different/similar tissue/cell types for EG interactions. For example, higher similarities are among blood cell lines, while other tissues displayed lower similarities with the other tissue/cell lines.

### Runtime of the EAGLE

EAGLE is faster than other tools. For predicting 388,090 candidate EG interactions in GM12878, EAGLE only took about only 2 minutes on Dell processor with 10 CPUs and memory of 32GB, while Ripple, TargetFinder, IM-PET, and JEME require ∼10, ∼50, ∼120, and ∼140 minutes, respectively.

## Discussion

Although several computational methods have been developed to predict EG interactions, these methods often require specific features (8–11). Therefore, these methods cannot be widely applied to many tissues, cell lines, or cell types. To predict the EG interactions in many tissue/cells, we developed a method that requires a small number of features, which are mainly derived from enhancer annotation and gene expression. Comparing with other tools, our method, EAGLE, has the following novelties: (1) The enhancers were integrated from multiple enhancer-related high through approaches, while the other tools (e.g. JEME, targetFinder, IM-PET, and RIPPLE) often predicted enhancers from a single technology (e.g. H3K27ac/H3K4me1 histone modification) (8–11, 24). We ensured the quality of enhancers. (2) We used six discriminative features, of which three (EWS, GWS, and WEEC) were never reported before. (3) The EAGLE model could be easily constructed in other species (e.g. mouse) and applied to predict EG interactions in other species.

GS, EWS, GWS, and WEEC are new features used for enhancer-target prediction. The cross-sample validation showed that they could greatly improve the performance, suggesting the usefulness of these new features (Fig S4). We also evaluated the effective importance of each feature by measuring the impact of permuting out-of-bag feature observation on the whole performance. These features have varying levels of contribution to the performance. Some features (e.g., H3K4me1) showed a limited difference between positives and negatives. However, the result indicated all six features have an effective contribution to the overall performance.

The distance between the enhancers and their potential targets is the most informative feature in the prediction. However, we are not able to separate the positives and negatives solely based on the feature. First, it is obvious that positive and negative pairs have a large overlap in distance distribution (Fig S3A). Second, if we use DIS as the only prediction feature, the area under ROC are 89.86% in self-test and 88.51% in cross-sample test. In contrast, if we include all the six features, the corresponding values are 95.65% in self-test and 93.38% in cross-sample test (Fig S3B and C). Third, the distances in positive EG pairs are still much larger than those between the enhancers and the nearest genes, suggesting that we cannot predict the EG pairs simply based on the nearest genes. Finally, like DIS, other features such as EGC, GS, and WEEC, could also reflect the tissue specificity of enhancer-gene interactions. In fact, we analyzed the importance of different features and found that DIS contributed 47% to the overall performance (Fig 5).

Among the six features, the enhancer-gene correlation (EGC) and enhancer-enhancer correlation (WEEC) were based on multiple cell types. However, they are still informative to predict tissue-specific interactions. For example, if the activity of one enhancer is correlated with one gene, it does not mean that the enhancer regulates the gene in all the cell types. The enhancer might only regulate the genes in the cell types where both the enhancer and the gene show high activity. The interactions do not occur in other cell types, although the information obtained from these cell types help us to establish the correlation.

We chose 1Mbp as the length of the scanned region because previous studies indicated that >99% of the real EG interactions were with a distance less than 1Mbp (8, 11). Our own analysis of ChIA-PET data also indicated that the number of genomic interactions decreases quickly with the increasing genomic distance (Fig S3D). Only 0.03% of genomic interactions were found to be from two regions with a distance greater than 1Mbp. We performed the EG interaction prediction with various cutoffs (Fig S3D). The number of positive interactions starts to saturate after 1Mbp, while the number of false positives keeps increasing. Indeed, similar scanned regions were used for many other prediction tools (e.g. IM-PET and JEME). Therefore, we believe that 1Mbp will cover the majority of EG interactions and has little impact on the predictions.

The two features EWS and GWS seem not very intuitive in this work. One important lesson we learned from enhancer biology is that enhancers are not necessary to regulate the nearest genes. There could be several other genes or enhancers located between the enhancers and their targets. We were interested in whether the number of genes (or enhancers), or the activity of these genes (or enhancers) could be informative features to predict enhancer-target relationships. After exploring different quantities, we found that EWS and GWS are useful in prediction. These two terms basically described the enhancer (or gene) activity normalized by the distance between an enhancer and target gene. In other words, if an enhancer interacts with a promoter, there are more active enhancers (or genes) between the interaction pair.

We could include more genomic features to improve the prediction. For example, we could include DNA binding motifs as additional features. However, it is a trade-off between adding more features and better prediction. If a program requires more features, it will become less flexible in practice because people often have limited datasets for a particular cell type. The selection of these six features is based on the availability of the datasets. For example, RNA-seq is widely used in labs and we expect people usually have the data available. We believe that more and more data types will become readily available and popular in the future. We will update the EAGLE by including more informative and easily accessible genomic features.

Multiple lines of evidence suggested that our prediction of enhancer-target interactions is reliable. We used the relevant histone modifications, ChIP-seq for TFs and eQTL enrichment to validate them. Unlike the non-interacting enhancers, the predicted interacting enhancers are significantly enriched for H3K27ac, H3K4me1 and H3K9ac modifications. Similarly, promoters in putative interactions also showed generally much higher signals than non-interacting ones in RAD21, CTCF, and H3K4me3 marks. We integrated genome-wide eQTL data across 48 tissues to detect the genotype-phenotype associations in DNA interactions and compared them with our predicted interactions.

The results showed that many predicted positives were supported by eQTLs, much higher than the predicted negatives. This result indicates that the enhancers interacting promoters have distinct properties than those do not interact with promoters.

We believe that the model captures the general rule for the interactions and the input data (e.g., gene expression and enhancer annotation) for each cell line contain the tissue specificity information. Therefore, even if our model was trained on a few cell types, the model can still be used to predict EG interactions in a variety of cell types. To demonstrate our point, we trained the models using cell lines of GM12878, MCF-7, and HeLa-S3, respectively (Fig. S5). We then predicted EG interactions in GM12878 using enhancers and expression data in GM12878 as input. The predicted EG interactions in GM12878 using these three models showed that each model predicted similar percent (around 11%) of positives overlapping with whole blood eQTLs and this percentage was much higher than that (∼7%) in other tissues, as well as that (around 0.7%) in negative controls (Fig. S10). These results indicated that the tissue-specific EG interactions were mainly achieved from the tissue-specific input data (e.g., enhancer annotation and gene expression), while the predicted model contained the general rules of EG interactions regardless of the cell line that was used to build the model.

In conclusion, we developed a common predictor requiring only the basic enhancer and gene information. With a simple input, our tool can be easily applied to predict interactions among new cis-regulatory genomic regions in new tissue/cells where not enough data are needed. The genome-wide predictions for human and mouse are available as a web resource at http://www.enhanceratlas.org/.

## Materials and methods

### Identification of enhancers and genes

Previous tools (e.g. JEME, TargetFinder, and RIPPLE) selected ChromHMM-predicted active enhancers by the chromatin state segmentation as the gold standard for training enhancers (9–11, 24). The enhancers defined by ChromHMM were based on histone modifications (24). Besides histone modification, many other high-throughput approaches (e.g. EP300, DHS, and CAGE) could also identify enhancers. To obtain reliable enhancers, we used ten independent high-throughput experimental tracks to identify the consensus enhancers by an unsupervised learning method (18). The high throughput approaches used to define enhancers include “TF-binding”, “DHS”, “Histone”, “EP300”, “POLR2A”, “CAGE”, “FARIE-seq”, “MNase”, “GRO-seq”, and “STARR”. Finally, we obtained 2,370,159 and 1,351,219 enhancers from all 110 and 89 tissue/cell types for human and mouse, respectively. We used the synthesized signal intensities from different genomic profiling as a proxy for the enhancer activity.

To estimate the gene expression values, we collected the RNA-seq from GEO datasets, UCSC genome browser and Roadmap data portal for human and mouse. Totally, 110 and 89 tissue/cells have RNA-seq data of good quality in human and mouse, respectively. For each gene, its promoter was defined as the TSS-containing regions 5kbp upstream and 0.5kbp downstream of the relative TSS based on a genomic position analysis of known promoters extracted from Broad ChromHMM resource (http://rohsdb.cmb.usc.edu/GBshape/cgi-bin/hgTrackUi?db=hg19&g=wgEnco35deBroadHmm).

### Definition and computation of the features

#### EGC

Recent studies showed that active enhancers were correlated with target gene expression patterns and this correlation could be used to infer their regulatory relationships (1, 14). Moreover, enhancer-promoter interactions also displayed specificity across different cell types (25). We utilized the signal values integrated from multiple high-throughput experimental tracks as enhancer activities and then build the correlation profiles between enhancer activity and gene expression levels across 110 tissue/cell types in human and 89 tissue/cell types in mouse, respectively. Given an enhancer *e* and a gene *g* across *m* tissue/cell types, the EGC could be defined as the Pearson Correlation Coefficient:

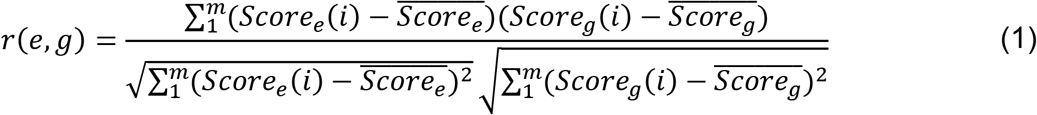

Where Score_e_(i) and Score_g_(i) represent the signal of enhancer and gene in i th tissue/cell type, respectively, while 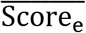 and 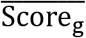 mean the average signal of enhancers and the average signal of genes across all tissue/cell types, respectively.

#### GS

For each tissue/cell, we define GS as the gene FPKM score in the processed RNA-seq data file. Since enhancers are the distal cis-regulatory elements that activate gene transcription, the gene scores in EG interactions should be generally higher than the ones in non-interacting pairs. By the empirical data in K562, the expression levels of genes in enhancer-target relationships are significantly different from the ones in non-interacting pairs (median values 4.743 vs. 0.289; p-value = 5.1e-31). Similarly, the difference in MCF-7 cells is also significant (median value 5.798 vs. 0.348; p-value = 2.0e-28).

#### DIS

This feature was defined as the genomic distance between the gene transcription start site and the enhancer.

#### EWS

Since only 7% of enhancer-promoter interaction loops selected the nearest gene for regulation (16), we expect that many active enhancers are located between the enhancer-target pair of interest. Assume that *m* enhancers located in the window between the enhancer and gene of one EG pair, and then EWS can be defined as:

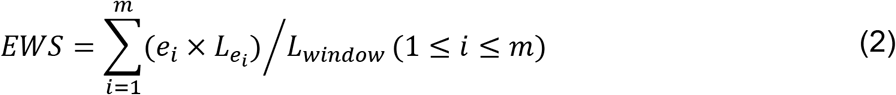

Where e_i_, 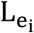 and L_window_ represent the average signal of enhancer *i*, the length of enhancer *i* and the length of the whole window, respectively. In K562, the value of EWS for positive pairs shows a significant difference from the negative ones (Median values 0.241 vs 0.049; p<2.2e-16).

#### GWS

Similar to EWS, we assume m genes located in the window and the gene window signal for the genes located between the enhancer and gene of one EG pair is defined as:

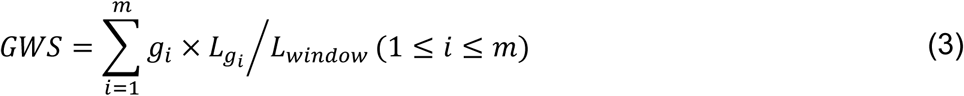

Where g_i_, 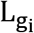, and L_window_ represent the signal of gene *i*, the length of gene *i* and the length of the whole window, respectively. The value of EWS for positives and negatives is significantly different in K562 (Median values 15.294 vs. 3.955; P<2.2e-16).

#### WEEC

Multiple enhancers often co-regulate one target gene. We expect the enhancer of one real EG pair should have a good correlation with other enhancers around the promoter of this pair. For m enhancers 1Mbp upstream or downstream one gene across n tissue/cell types, the correlations of the *i*th enhancer (1 ≤ *i* ≤ m) with the other m-1 enhancers were calculated. Then the WEEC for this enhancer is defined as:

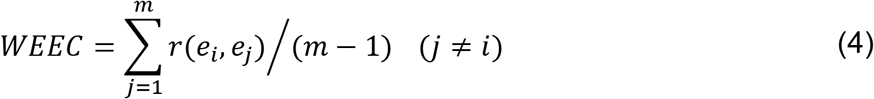

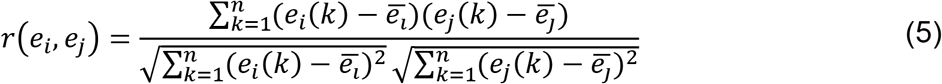

Where e_i_(*k*) and e_j_(*k*) represent the signals of the *i*th and the *j*th enhancers in the *k*th tissue/cell type, respectively, while e̅_i_ and e̅_j_ mean the average signals of the ith and the jth enhancers across all *n* tissue/cell types.

### Preparing of training datasets

We mapped all the enhancers to the regions 1 Mbp upstream/downstream the genes and integrated all the candidate EG pairs within these regions. For the human, the ChIA-PET data marked by anti-RNA polymerase II antibody were used as a gold standard to define the positive pairs. Since no ChIA-PET data of good quality in mouse, we choose the Hi-C data with a high resolution of 2.5kb to build the training datasets (26). Generally, the training EG pairs are selected with three criteria: (i) The enhancers are supported by at least 50% of the high-throughput experimental evidence (e.g. P300, DNase, TF-Binding, CAGE, and Histone). (ii) The potential target genes are expressed. No matter in positive or negative pairs, all the genes are assigned with a RNA-seq expression signal (FPKM value>0). (iii) The pair is overlapped (positives) or not overlapped (negatives) with ChIA-PET or Hi-C.

### Model training

We tried several learning methods (SVM, KNN, Discriminant, Decision tree, and ensemble boosting) for training. Of all methods, ensemble boosting algorithm “AdaBoost” showed the best performance by 10-fold cross-validation (Fig S1). “AdaBoost” fits a series of weak classifiers that are slightly better than random ones and converts weak classifier to strong classifier by increasing or decreasing the weight of samples (Fig S11). Our EAGLE model randomly selected half of the positive EG pairs and the same number of negative EG pairs for training, using 50 decision trees by 30 learning cycles. We calculated the final weight of each pair by the classifier to determine if it is positive or not.

### Comparing with four existing EG prediction tools

In order to reasonably compare EAGLE with the other four tools, we used the same data for them. Since all of the existing tools used the GM12878 data for testing, we made a comparison on this cell line. Reliable enhancers are defined with multiple tracks by our previous method (18). We used the FPKM values from processed RNA-seq data to define gene scores. All candidate EG interactions are constructed by assigning all the enhancers 1Mbp upstream or downstream the center gene and used the ChIA-PET data as the gold standard to define the positives and negatives.

IM-PET was downloaded at http://tanlab4generegulation.org/IM-PET.html and was implemented in Linux platform. We used the same enhancer and gene profiles in GM12878 as input to predict the interacting pairs by both EAGLE (Trained by K562 ChIA-PET) and IM-PET. Then their relative pairs are analyzed to calculate the relative performance. No function in IM-PET was used for self-testing, so only the across-sample testing is used in this comparison. We also performed this comparison based on MCF-7 and Hela-S3 cell lines.

RIPPLE was downloaded at http://pages.discovery.wisc.edu/~sroy/ripple/download.html was run by Matlab. To enable our GM12878 data to be predicted by RIPPLE, we integrated many features (e.g. Dnase1, H3k27ac, H3K4me3) the tool required from http://ftp.ebi.ac.uk/pub/databases/ensembl/encode/integration_data_jan2011/byDataType/peaks/jan2011/histone_macs/optimal/ and http://ftp.ebi.ac.uk/pub/databases/ensembl/encode/integration_data_jan2011/byDataType/peaks/jan2011/spp/optimal. For MCF-7 and Hela-S3, we collect the corresponding data from GEO datasets.

TargetFinder was downloaded at https://github.com/shwhalen/targetfinder and performed with python in window platform. We collected and integrated as many as possible features for each GM12878 pair by the raw peak files in TargetFinder. To make TargetFinder model consistent with EAGLE, we also used the K562 data in TargetFinder for training and took the trained model to predict the interacting pairs in the same GM12878 data. Specifically, the common features in both K562 and GM12878 were used for training and testing. In the same way, we integrated the data in MCF-7 and Hela-S3 for testing.

JEME was downloaded at https://github.com/yiplabcuhk/JEME/ and implemented by sh and R languages in Linux platform. We used the same GM12878, MCF-7 and Hela-S3 pairs for JEME and EAGLE. To make the data be predicted by JEME, we collected and curated four features according to the format of JEME “Roadmap” model required. The predicted pairs with low or high scores are used for comparison.

### Importance of training features

For six features in each cell line, we used the Matlab function “oobPermutedPredictorImportance” to estimate the feature importance by permutation of out-of-bag feature observations.

### Validation with genomic features and eQTL

The data of genomic features (H3K27ac, H3K4me1, H3K9ac, CTCF, RAD21, and H3K4me3) were downloaded from the website (http://ftp.ebi.ac.uk/pub/databases/ensembl/encode/integration_data_jan2011/byDataType/signal/jan2011/bedgraph/). The density analysis and heatmap for enhancers and promoters were based on the 2.5 kb window around the center of the enhancers and TSS of genes, respectively. For eQTL, we used the latest data named “GTEx Analysis V7” (https://gtexportal.org/home/datasets). In the GTEx database, q-value was used to decide the genes significantly associated with the genetic variance (21).

### Software implementation

EAGLE was implemented in Perl and Matlab with learning ensemble methods. All the codes are put in the GitHub website https://github.com/EvansGao/EAGLE.

## Acknowledgments

We thank Drs. Minwen Hu, Jianbo Pan, and Jie Wang for their insightful comments.

## Supporting information captions

**S1 Fig. Comparison of different machine learning approaches.** 71118 pairs with 35559 positives and 35559 negatives in K562 were taken as the common training data. 10-fold cross validation was used for all approaches. The functions “fitcensemble”, “fitctree”, “fitcdiscr”, “fitcknn” and “fitcsvm” in matlab, and “lm” in R were adopted to EAGLE, Decision tree, Discriminant, KNN, SVM, and linear regression respectively.

**S2 Fig. Six differentiable features in GM12878.** (A) Enhancer activity and gene expression profile correlation (EGC). (B) Gene score (GS) from the RNA-seq data. (C) Distance (DIS) between enhancer and gene in a pair. (D) Enhancer window signal (EWS) measuring the mean enhancer signal in the region between enhancer and promoter (E) Gene window signal (GWS) evaluating the mean gene expression level in the region between enhancer and promoter (F) The weight of enhancer-enhancer correlation (WEEC). The positive, negative and random enhancer-gene pairs were obtained from ChIA-PET dataset in K562. The *P* values were calculated using Student *t* test.

**S3 Fig. Distribution of enhancer-gene distances in positives, negatives and pairs with nearest genes and individual performance of DIS.** (A) Distributions of distances in positives and negatives of K562. (B) Individual self-test performance of DIS and other features in K562. (C) Individual cross-sample test performance of DIS and other features with training in K562 and testing in GM12878. (D) Changes of the number of positives/negatives and the prediction performances with various cutoffs of scanned regions. (E) Comparison of distances between positives (Marked as “Real”) and pairs with nearest genes in K562.

**S4 Fig. Cross sample validation.** We trained the model using K562 and tested the model in GM12878. (A) Testing based on balanced data with 9732 positives and 9732 negatives in GM12878. Left panel is the ROC and right panel is the PR curves. (B) Testing using unbalanced data with 9732 positives and 48661 negatives in GM12878. Left panel is the ROC and right panel is the PR curves. We successively added the features (EGC, GS, EWS, GWS, EEC and DIS) to show the improving performance.

**S5 Fig. The performances of prediction models constructed in other three cell lines.**

**S6 Fig. Cross-sample validation of performances for enhancer-gene prediction tools in other cell lines.**(A) Relative AUROCs and AUPRs of all tools in MCF-7 (B) AUROCs and AUPRs of five tools in Hela-S3. The cross-sample validation was performed with the training in K562 and testing in other cell lines (see Methods).

**S7 Fig. Evaluation of feature importance using self-testing.** (A) Performances (AUROC and AUPR) gradually improved with successive adding of the training features in K562. (B) Performance (AUROC and AUPR) increasing with adding the training features one by one in MCF-7. For each cell line, the self-testing used one half of the data for training and the other half for testing.

**S8 Fig. The features in mouse lung.** (A) Enhancer activity and gene expression profile correlation (EGC) (B) Gene signal from the RNA-seq data. (C) Distance between enhancer and gene in a pair. (D) Enhancer window signal measuring the mean enhancer signal in the region between enhancer and promoter (E) Gene window signal evaluating the mean gene expression level in the region between enhancer and promoter. The P values were calculated by the Student t test.

**S9 Fig. Self-testing and cross-sample test with lung model in mouse.** (A) Self-testing by PR plot in lung. (B) cross-sample test on spleen with PR plot by lung model.

**S10 Fig. The correlation between eQTLs and predicted EG interactions by different prediction models.** The enhancers and expression data in GM12878 were taken as the input. (A) The similar percent (around 11%) of positives and percent (around 0.7%) of negatives in the predicted EG interactions of GM12878 by different models, overlapping with eQTLs in whole blood. (B) The simimar percent (around 11%) of positives overlapping with whole blood eQTLs much higher than that (∼7%) in other 47 tissues.

**S11 Fig. The overview of ensemble boosting algorithm training process.** (A) Weak classifier is set to classify all enhancer-gene interaction sites assigned with equal weights in the initial stage. (B)The subsequent classifier keeps track of previous classifier’s errors and starts to distinguish the positives from negatives by randomly increasing positive sites’ weights or decreasing negatives’ weights. (C) With utilizing more and more success of previous classifiers, the new generated classifier is trained with a good classification on most sites. (D) The classifier becomes perfect when all sites’ weights are appropriately changed. Generally speaking, the boosting algorithm made each classifier trained with taking into account the previous one’s success. In each step of training, the weights of some sites will be redistributed. Specially, misclassified sites will change its weights to emphasize their difficulties. Then subsequent new classifiers will focus on them during the new training.

**S1 Table. Summary of datasets collected for mouse enhancers in 156 tissue/cell types.** Each tissue/cell type has at least three tracks and each enhancer is supported by at least one half of the tracks in the relative tissue/cell type.

