## Supplemental figures for "EAGLE: an algorithm that utilizes a small number of genomic features to predict tissue/cell type-specific enhancer-gene interactions"

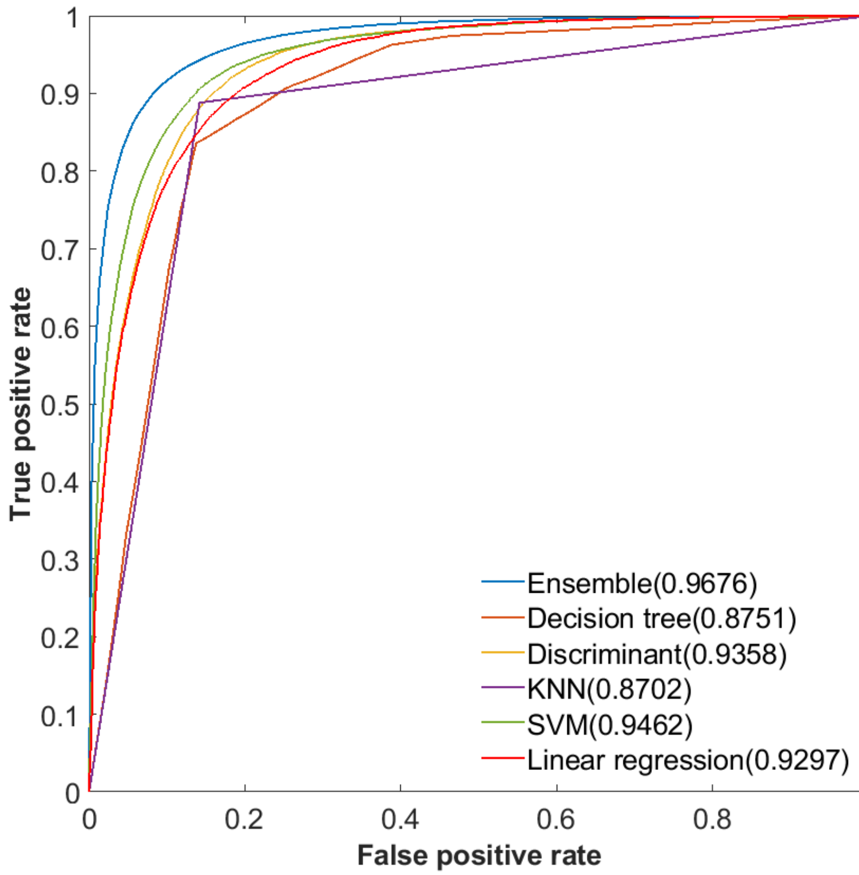


**Figure S1.** Comparison of different machine learning approaches. 71118 pairs with 35559 positives and 35559 negatives in K562 were taken as the common training data. 10-fold cross validation was used for all approaches. The functions “fitcensemble”, “fitctree”, “fitcdiscr”, “fitcknn” and “fitcsvm” in matlab, and “lm” in R were adopted to EAGLE, Decision tree, Discriminant, KNN, SVM, and linear regression respectively.

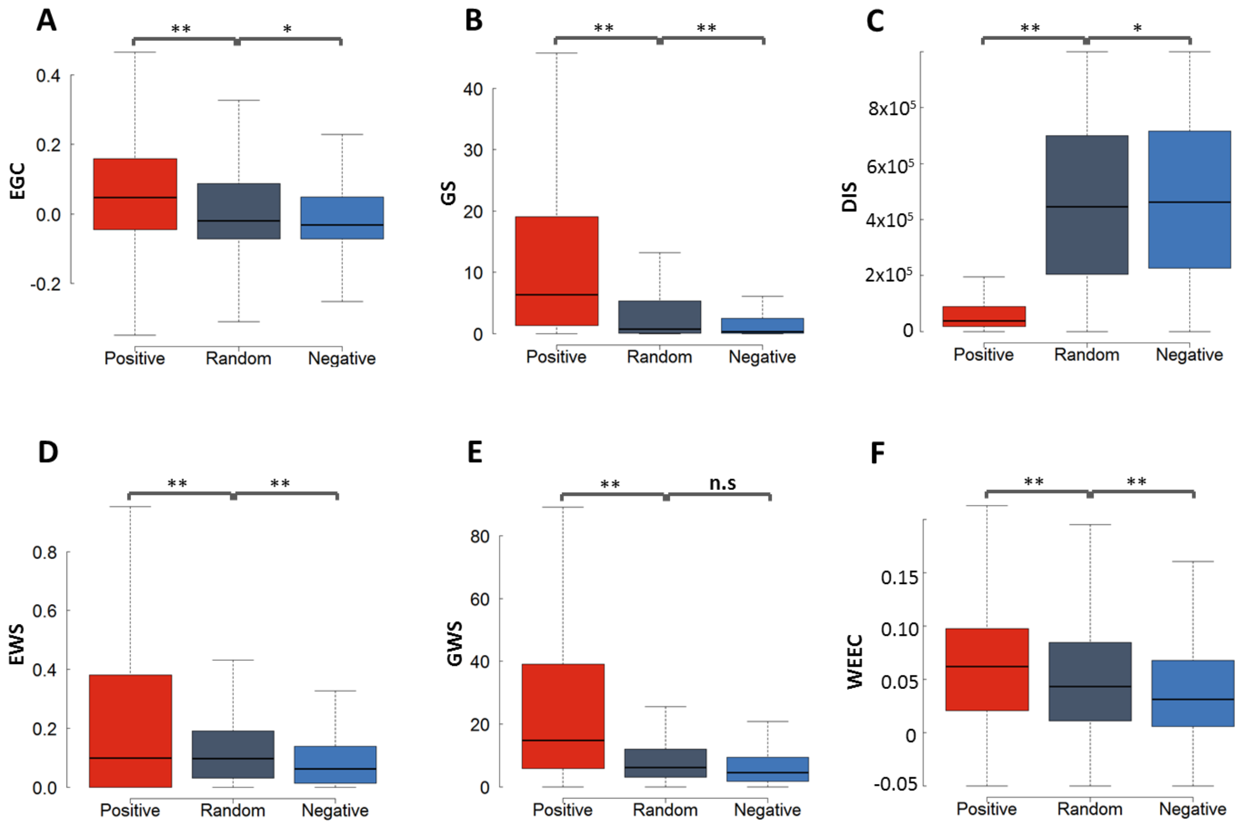


**Figure S2.** Six differentiable features in GM12878. (A) Enhancer activity and gene expression profile correlation (EGC). (B) Gene score (GS) from the RNA-seq data. (C) Distance (DIS) between enhancer and gene in a pair. (D) Enhancer window signal (EWS) measuring the mean enhancer signal in the region between enhancer and promoter (E) Gene window signal (GWS) evaluating the mean gene expression level in the region between enhancer and promoter (F) The weight of enhancer-enhancer correlation (WEEC). The positive, negative and random enhancer-gene pais were obtained from ChIA-PET dataset in K562. The *P* values were calculated using Student *t* test.

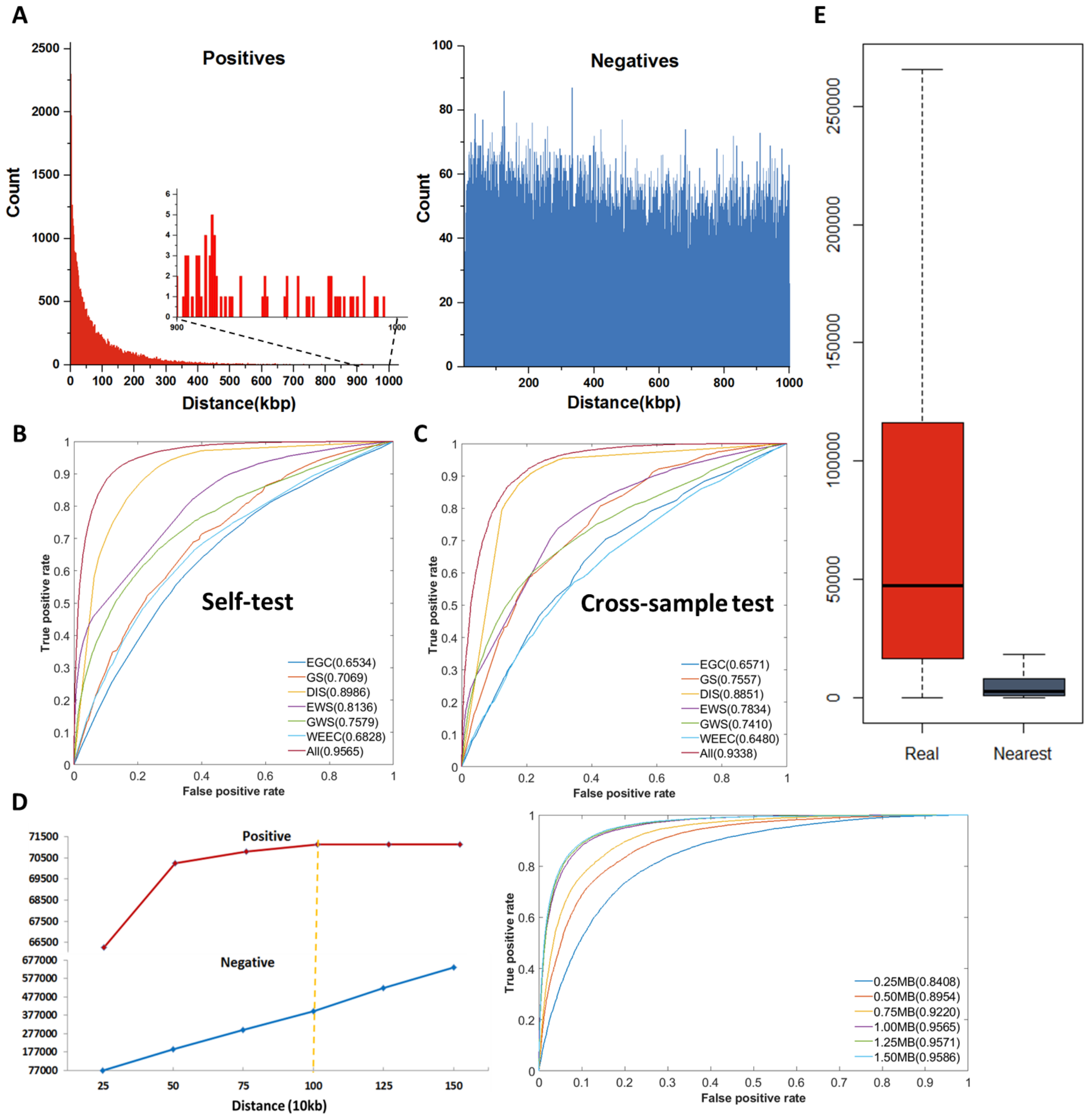


**Figure S3.** Distribution of enhancer-gene distances in positives, negatives and pairs with nearest genes and individual performance of DIS. (A) Distributions of distances in positives and negatives of K562. (B) Individual self-test performance of DIS and other features in K562. (C) Individual cross-sample test performance of DIS and other features with training in K562 and testing in GM12878. (D) Changes of the number of positives/negatives and the prediction performances with various cutoffs of scanned regions. (E) Comparison of distances between positives (Marked as “Real”) and pairs with nearest genes in K562.

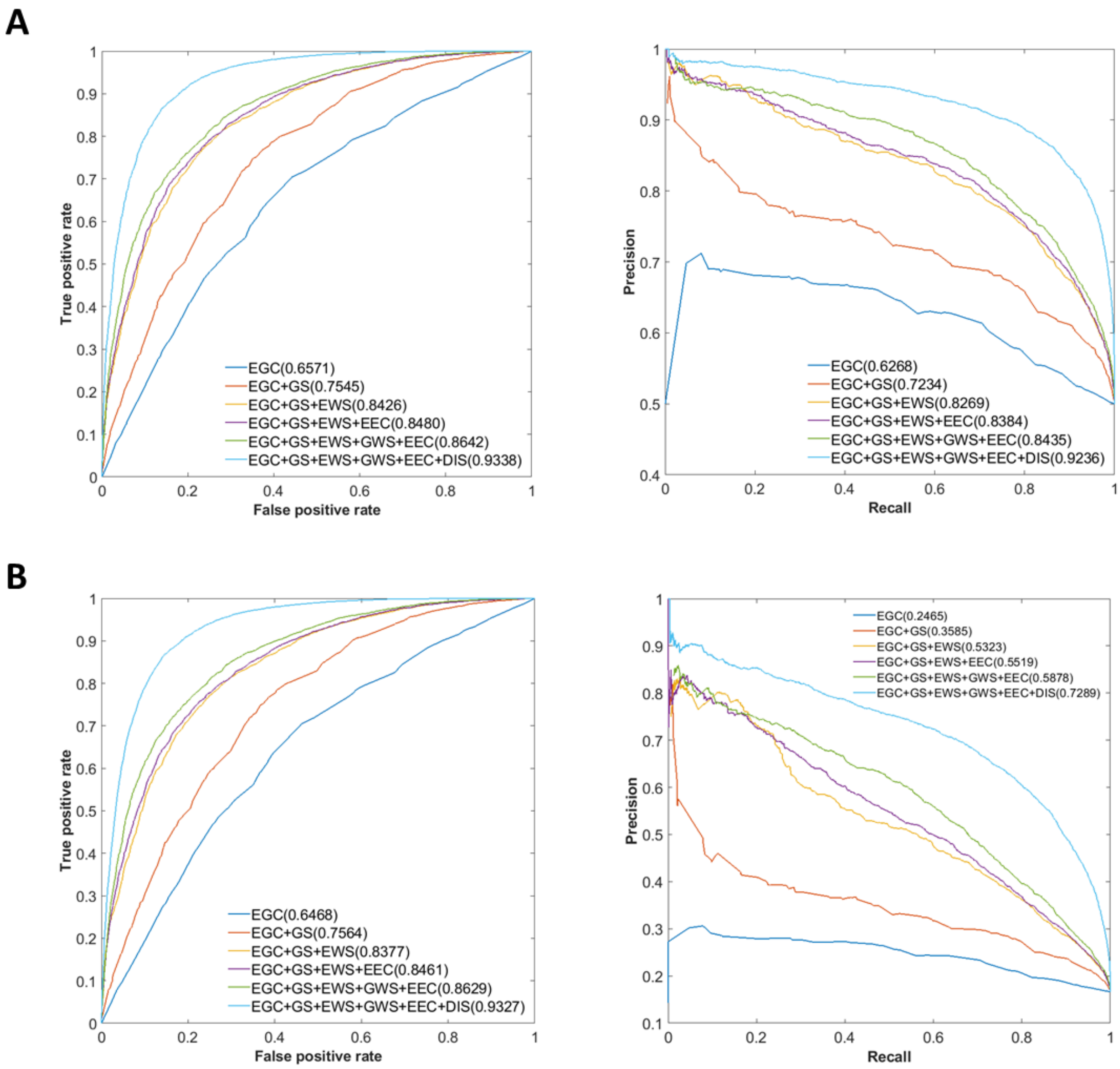


**Figure S4.** Cross sample validation. We trained the model using K562 and tested the model in GM12878. (A) Testing based on balanced data with 9732 positives and 9732 negatives in GM12878. Left panel is the ROC and right panel is the PR curves. (B) Testing using unbalanced data with 9732 positives and 48661 negatives in GM12878. Left panel is the ROC and right panel is the PR curves. We successively added the features (EGC, GS, EWS, GWS, EEC and DIS) to show the improving performance.

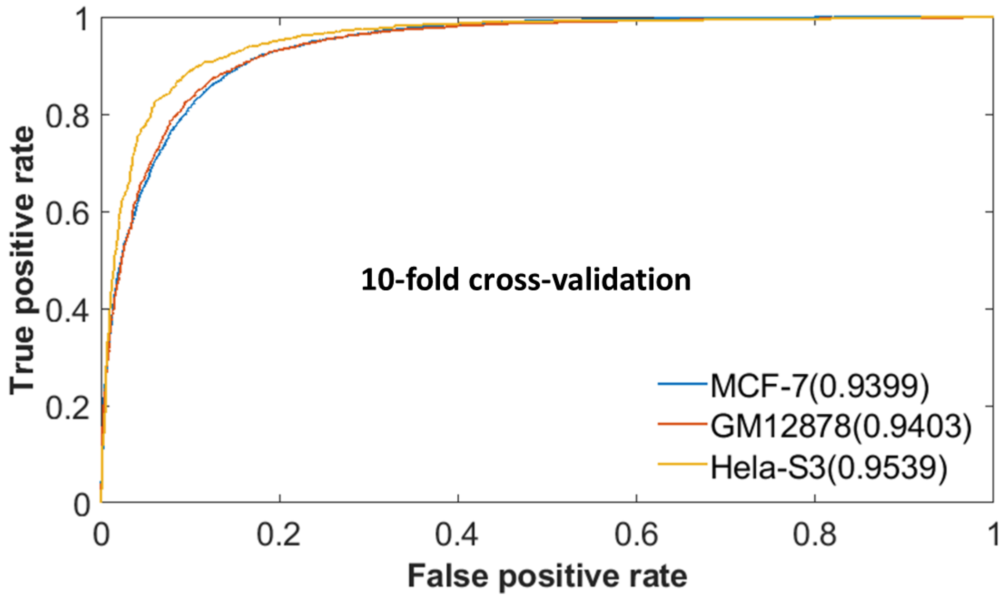


**Figure S5.** The performances of prediction models constructed in other three cell lines.

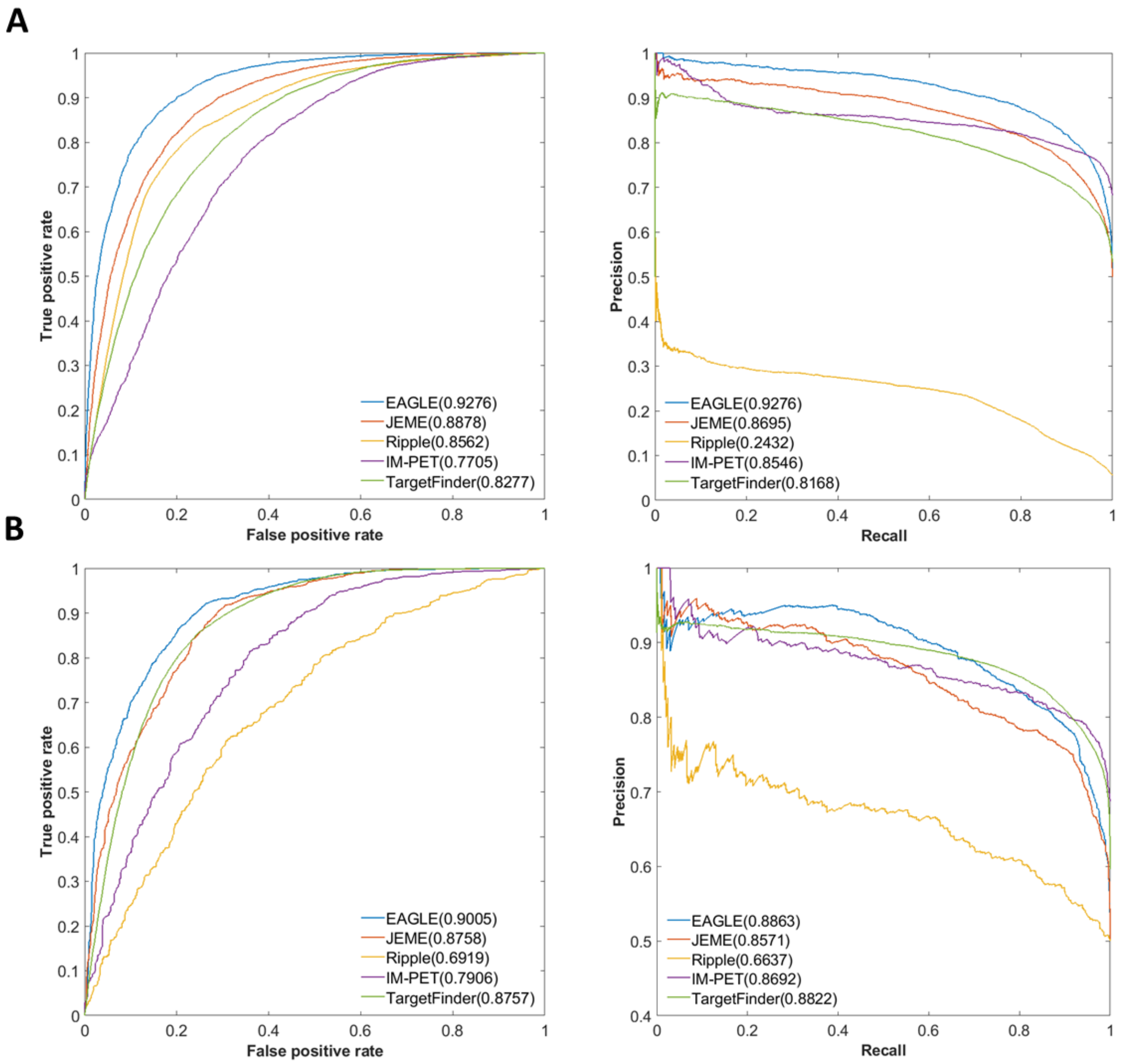


**Figure S6** Cross-sample validation of performances for enhancer-gene prediction tools in other cell lines. (A) Relative AUROCs and AUPRs of all tools in MCF-7. (B) AUROCs and AUPRs of five tools in Hela-S3. The cross-sample validation was performed with the training in K562 and testing in other cell lines (see Methods).

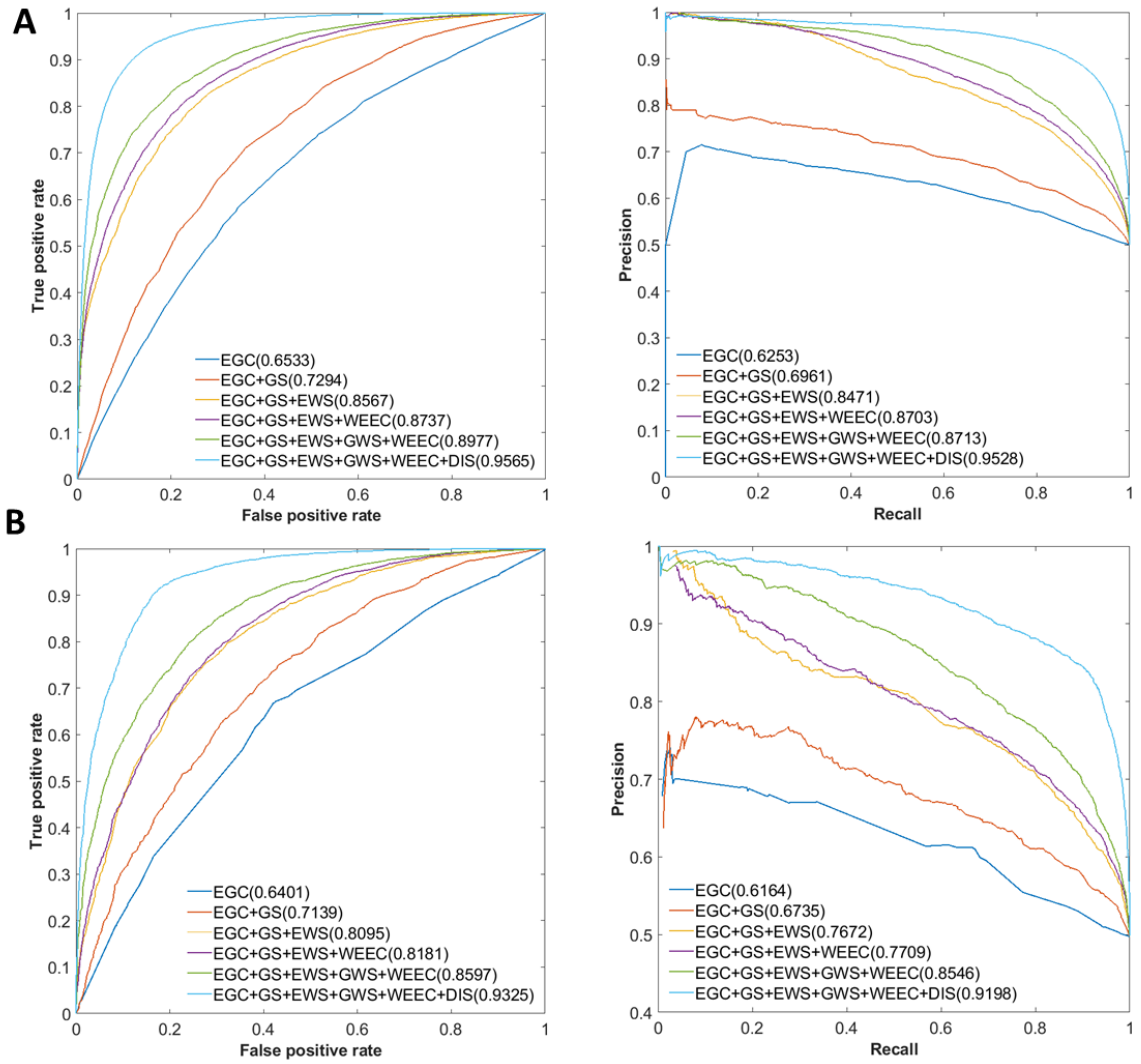


**Figure S7.** Evaluation of feature importance using self-testing (A) Performances (AUROC and AUPR) gradually improved with successive adding of the training features in K562. (B) Performance (AUROC and AUPR) increasing with adding the training features one by one in MCF-7. For each cell line, the self-testing used one half of the data for training and the other half for testing.

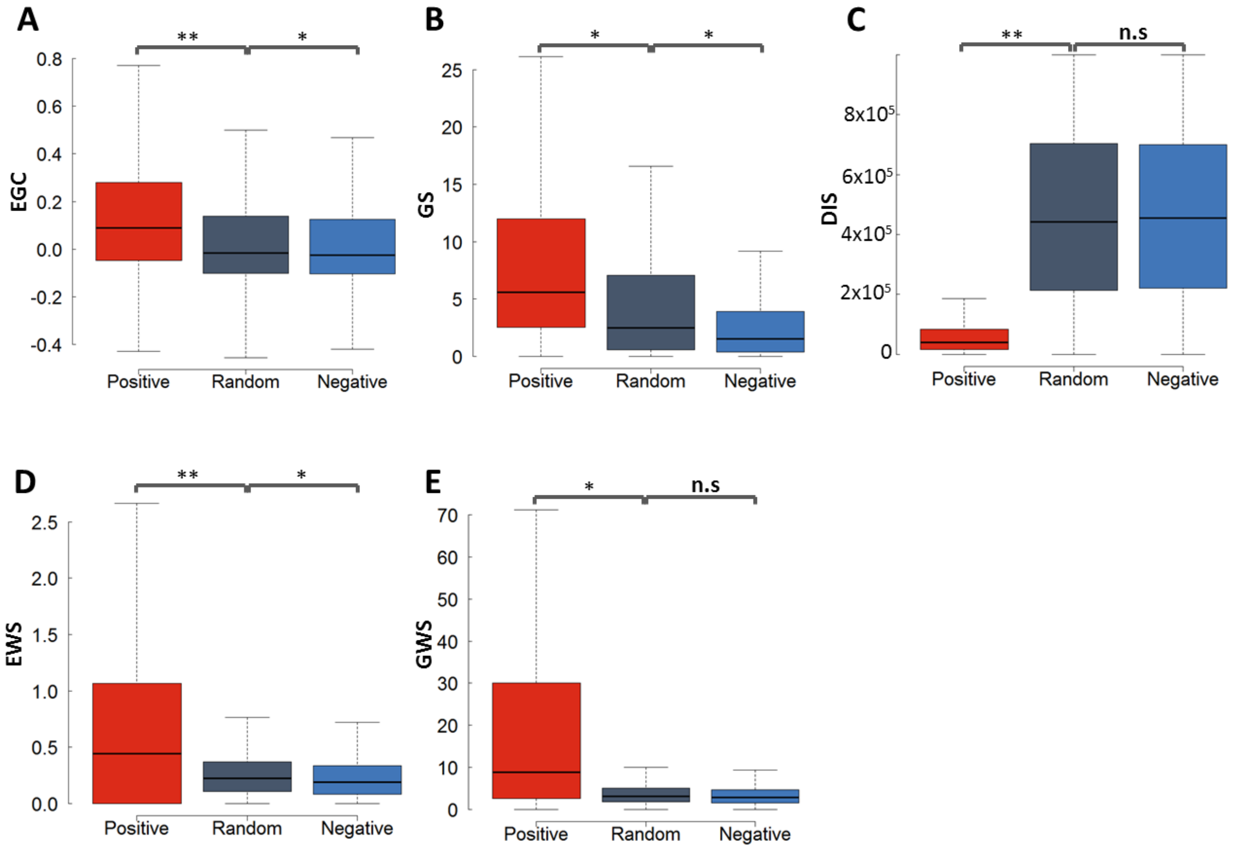


**Figure S8.**The features in mouse lung. (A) Enhancer activity and gene expression profile correlation (EGC) (B) Gene signal from the RNA-seq data. (C) Distance between enhancer and gene in a pair. (D) Enhancer window signal measuring the mean enhancer signal in the region between enhancer and promoter (E) Gene window signal evaluating the mean gene expression level in the region between enhancer and promoter. The *P* values were calculated by the Student t test.

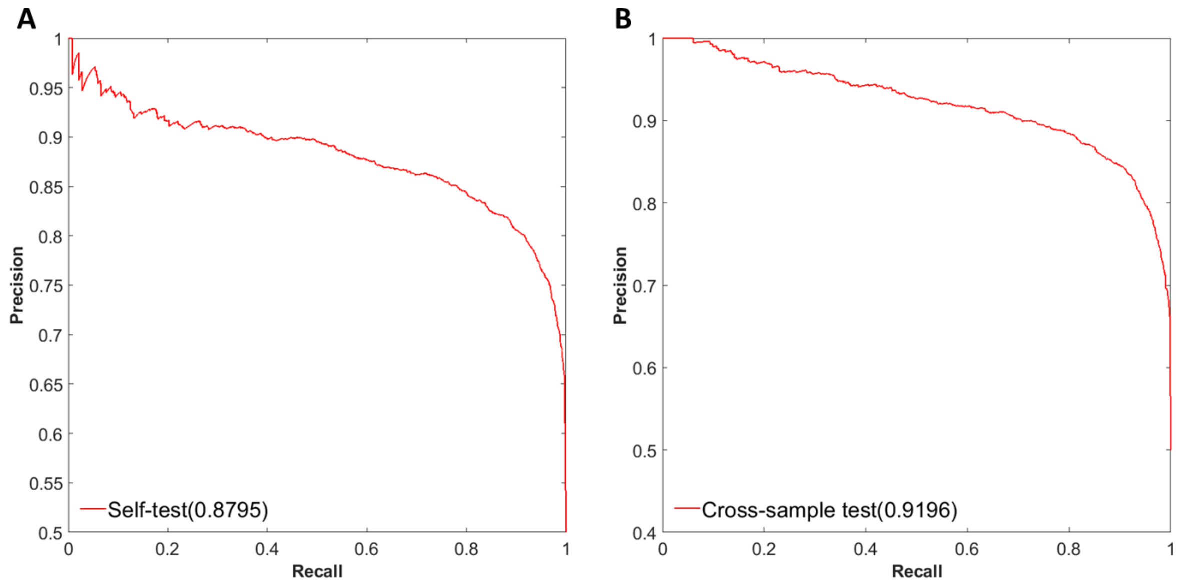


**Figure S9** Self-testing and cross-sample test with lung model in mouse. (A) Self-testing by PR plot in lung. (B) cross-sample test on spleen with PR plot by lung model.

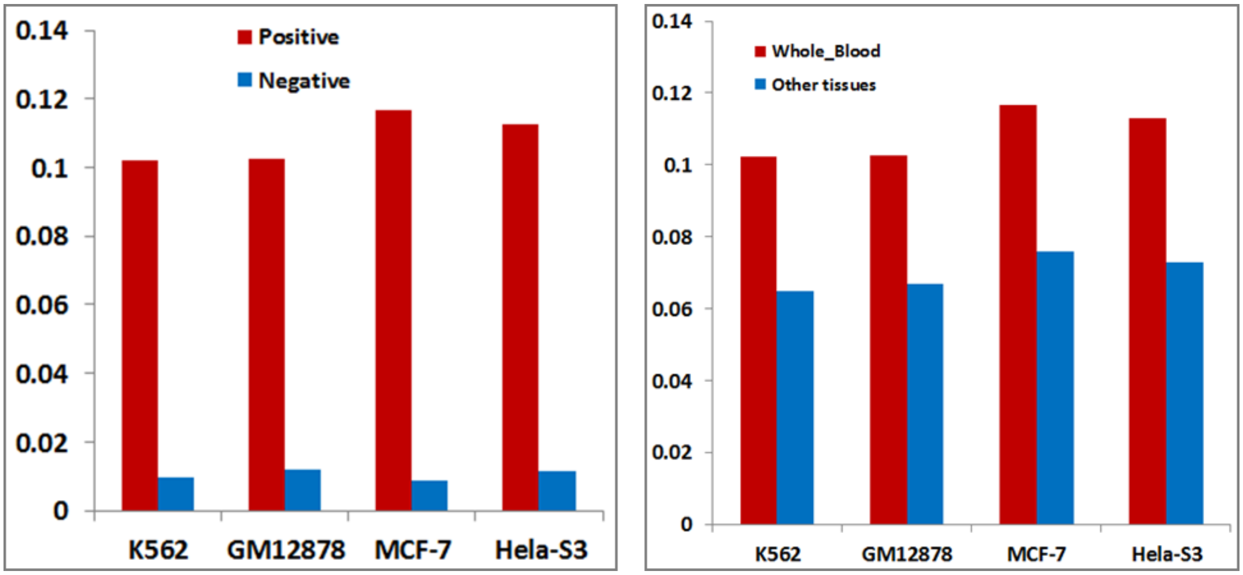


**Figure S10.** The correlation between eQTLs and predicted EG interactions by different prediction models. The enhancers and expression data in GM12878 were taken as the input. (A) The similar percent (around 11%) of positives and percent (around 0.7%) of negatives in the predicted EG interactions of GM12878 by different models, overlapping with eQTLs in whole blood. (B) The simimar percent (around 11%) of positives overlapping with whole blood eQTLs much higher than that (~7%) in other 47 tissues.

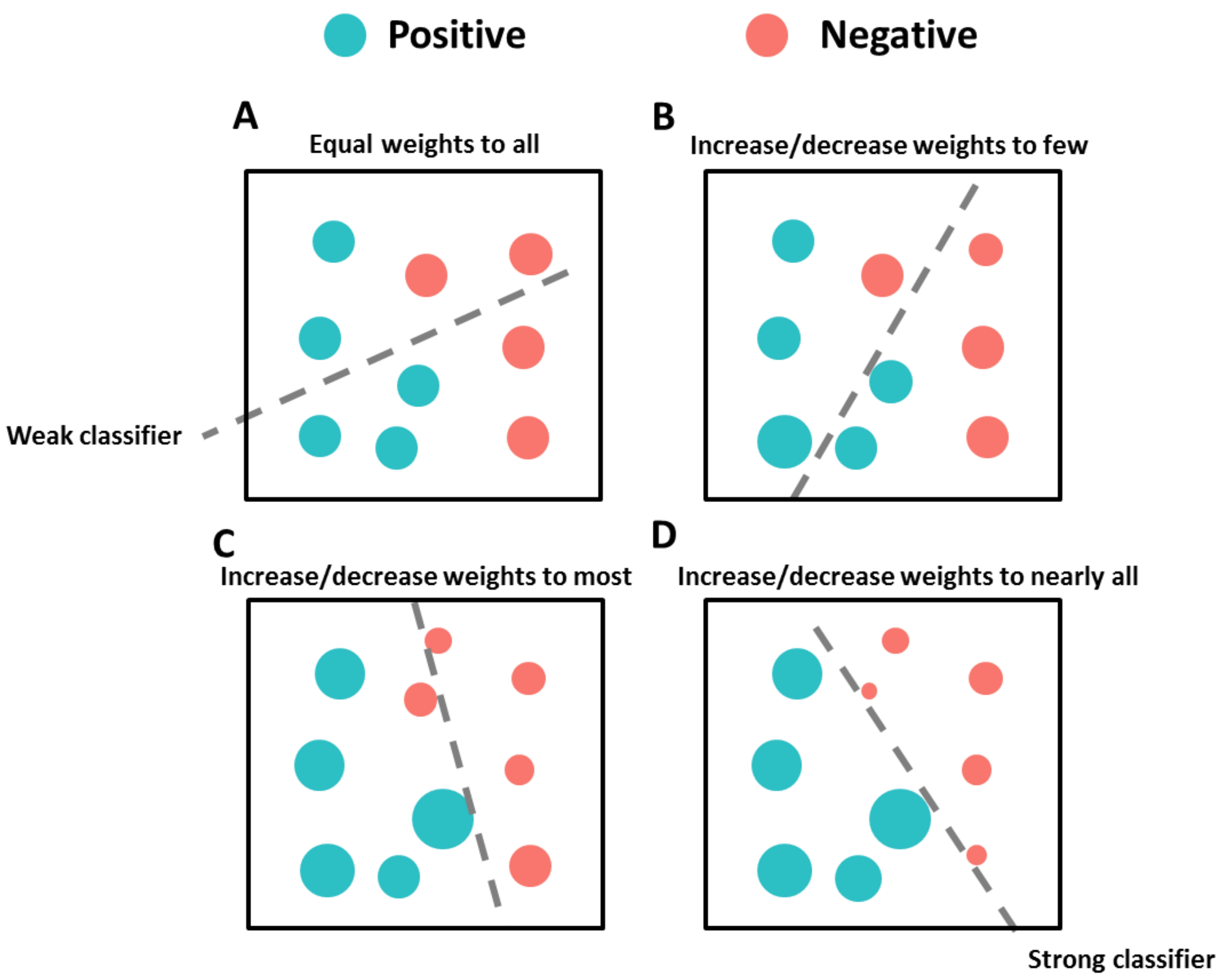


**Figure S11** The overview of ensemble boosting algorithm training process. (A) Weak classifier is set to classify all enhancer-gene interaction sites assigned with equal weights in the initial stage. (B)The subsequent classifier keeps track of previous classifier's errors and starts to distinguish the positives from negatives by randomly increasing positive sites' weights or decreasing negatives' weights. (C) With utilizing more and more success of previous classifiers, the new generated classifier is trained with a good classification on most sites. (D) The classifier becomes perfect when all sites' weights are appropriately changed. Generally speaking, the boosting algorithm made each classifier trained with taking into account the previous one's success. In each step of training, the weights of some sites will be redistributed. Specially, misclassified sites will change its weights to emphasize their difficulties. Then subsequent new classifiers will focus on them during the new training.
