## Supplemental table for "EAGLE: an algorithm that utilizes a small number of genomic features to predict tissue/cell type-specific enhancer-gene interactions"

### Supplementary Tables:

**Table S1**: Summary of datasets collected for mouse enhancers in 156 tissue/cells

| Tissue/cell | Approaches | Num. of datasets |
|---|---|---|
| B134 | TF-Binding;POL2;DHS;EP300 | 54 |
| BT3-L1 | TF-Binding;POL2;Histone;DHS;EP300;GRO-seq | 216 |
| 416B | Histone;TF-Binding;DHS | 15 |
| Astrocyte | TF-Binding;FAIRE;CAGE | 15 |
| AtT-20 | TF-Binding;EP300;Histone;GRO-seq | 15 |
| B220+ | TF-Binding;POL2;Histone | 18 |
| BAT | Histone;FAIRE;TF-Binding;POL2 | 31 |
| Bone_marrow | Histone;POL2;TF-Binding | 20 |
| Brain | Histone;TF-Binding;POL2;EP300;DHS | 32 |
| Brain_E14.5 | TF-Binding;DHS;POL2;Histone | 18 |
| Brown_preadipocyte_E18.5 | TF-Binding;Histone;POL2 | 63 |
| B_cell | TF-Binding;Histone;DHS;POL2 | 109 |
| B_cell_BM | Histone;DHS;TF-Binding | 32 |
| B_cell_lymph_node | TF-Binding;POL2;Histone;DHS | 33 |
| B_cell_spleen | TF-Binding;Histone;POL2;DHS;GRO-seq | 124 |
| C10 | Histone;TF-Binding;EP300;MNase | 39 |
| C3H10Thalf | TF-Binding;Histone;POL2 | 27 |
| Cardiomyocyte | TF-Binding;Histone;POL2;CAGE | 13 |
| CD172+DC | TF-Binding;EP300;Histone | 4 |
| CD19+ | TF-Binding;Histone;DHS;CAGE | 9 |
| CD24+DC | TF-Binding;EP300;Histone | 5 |
| CD4+ | Histone;TF-Binding;DHS;POL2;CAGE | 205 |
| CD4+CD8+ | DHS;TF-Binding;Histone;MNase;POL2 | 57 |
| CD4+Treg | TF-Binding;EP300;Histone | 21 |
| CD4-CD8- | DHS;Histone;TF-Binding | 10 |
| CD43- | POL2;DHS;TF-Binding | 32 |
| CD8+ | TF-Binding;EP300;Histone;DHS;POL2;CAGE | 80 |
| Cerebellum | TF-Binding;Histone;DHS;POL2;CAGE | 35 |
| Cerebellum_neonate | TF-Binding;DHS;Histone;POL2;CAGE | 41 |
| CH12 | DHS;POL2;TF-Binding;EP300;Histone | 40 |
| CMP | Histone;DHS;CAGE | 3 |
| Cortex | TF-Binding;POL2;Histone;CAGE | 17 |
| Cortex_embryo | TF-Binding;Histone;EP300 | 29 |
| Dendritic_cell | Histone;TF-Binding;POL2 | 30 |
| DR_cell | Histone;EP300;TF-Binding;POL2 | 6 |
| E14 | TF-Binding;Histone;MNase;DHS;FAIRE;POL2;EP300 | 276 |
| Embryoid_body_cell | TF-Binding;POL2;Histone | 40 |
| Embryo_body | DHS;Histone;CAGE | 18 |
| Epididymis | TF-Binding;Histone;CAGE | 18 |
| EpiLC | TF-Binding;Histone;EP300;POL2;FAIRE | 66 |
| EpiSC | TF-Binding;Histone;EP300;POL2;FAIRE | 21 |
| Erythroblast_fetal_liver | TF-Binding;Histone;POL2 | 11 |
| Erythroid_fetal_liver | TF-Binding;Histone;POL2 | 11 |
| Erythroid_spleen | TF-Binding;Histone;POL2;DHS | 6 |
| ESC_46C | TF-Binding;Histone;MNase;FAIRE;POL2;DHS | 53 |
| ESC_Bruce4 | Histone;POL2;EP300 | 13 |
| ESC_C6 | TF-Binding;Histone;POL2 | 10 |
| ESC_D0 | TF-Binding;Histone;POL2 | 22 |
| ESC_D3 | TF-Binding;Histone;POL2 | 34 |
| ESC_hemangioblast | Histone;TF-Binding;DHS | 7 |
| ESC_hematopoietic_progenitor | Histone;TF-Binding;DHS | 12 |
| ESC_hemogenic_endothelium | Histone;TF-Binding;DHS | 9 |
| ESC_J1 | Histone;DHS;TF-Binding;MNase;POL2 | 126 |
| ESC_KH2 | TF-Binding;POL2;GRO-seq | 96 |
| ESC_NPC | TF-Binding;FAIRE;POL2;MNase;Histone | 60 |
| ESC_R | Histone;EP300;TF-Binding;POL2 | 40 |
| ESC_R1 | DHS;TF-Binding;MNase;EP300;Histone;FAIRE | 77 |
| Fibroblast | Histone;TF-Binding;DHS | 9 |
| Forebrain_E11.5 | Histone;TF-Binding;EP300 | 18 |
| Forebrain_E12.5 | Histone;TF-Binding;POL2 | 19 |
| Forelimb_bud_embryo | TF-Binding;DHS;Histone | 14 |
| Forelimb_E11 | Histone;TF-Binding;CAGE | 4 |
| Forelimb_E13 | Histone;POL2;TF-Binding;CAGE | 12 |
| G1E | TF-Binding;Histone;DHS;POL2 | 42 |
| G1E-ER4 | Histone;POL2;TF-Binding;DHS | 113 |
| Gastrocnemius_muscle_myoblast | TF-Binding;Histone;DHS | 18 |
| Germline_stem_cell | POL2;TF-Binding;MNase | 9 |
| GMP | TF-Binding;DHS;Histone;CAGE | 11 |
| Heart | TF-Binding;DHS;Histone;POL2;EP300;FAIRE;GRO-seq | 74 |
| Heart_E11.5 | Histone;TF-Binding;EP300;CAGE | 29 |
| Heart_E12.5 | TF-Binding;Histone;POL2;CAGE | 28 |
| Heart_E14.5 | Histone;POL2;CAGE | 16 |
| Heart_neonate | DHS;Histone;EP300;CAGE | 15 |
| Hepatocyte | TF-Binding;EP300;CAGE | 13 |
| HFSC | DHS;Histone;TF-Binding | 20 |
| Hindbrain_E11.5 | DHS;Histone;TF-Binding | 14 |
| Hindlimb_bud_E11.5 | DHS;TF-Binding;Histone | 7 |
| HL-1 | Histone;EP300;TF-Binding | 12 |
| HPC7 | EP300;TF-Binding;Histone | 26 |
| HSC_BM | DHS;Histone;CAGE | 12 |
| Intestine | Histone;POL2;CAGE | 4 |
| Intestine_crypt_progenitor | Histone;DHS;TF-Binding | 5 |
| Intestine_embryo | Histone;TF-Binding;CAGE | 20 |
| IPSC | Histone;TF-Binding;POL2;DHS;MNase | 74 |
| Kidney | Histone;DHS;TF-Binding;POL2;EP300 | 26 |
| Kidney_embryo | Histone;TF-Binding;CAGE | 34 |
| Kidney_neonate | DHS;Histone;CAGE | 8 |
| Large_intestine_epithelial | Histone;TF-Binding;DHS | 33 |
| Lens_P1 | TF-Binding;Histone;POL2;FAIRE | 8 |
| Limb_E11.5 | DHS;Histone;TF-Binding;EP300 | 15 |
| Limb_E14.5 | DHS;Histone;POL2 | 17 |
| Liver_E14.5 | Histone;DHS;POL2;CAGE | 17 |
| Liver_neonate | DHS;Histone;TF-Binding;POL2;CAGE | 30 |
| Liver_tumor | TF-Binding;POL2;Histone | 23 |
| LSK | TF-Binding;DHS;Histone;CAGE | 24 |
| Lung | Histone;TF-Binding;DHS;POL2;EP300;CAGE | 20 |
| Lung_E14.5 | DHS;Histone;CAGE | 13 |
| Lung_neonate | DHS;Histone;CAGE | 8 |
| Lung_tumor | TF-Binding;DHS;Histone | 13 |
| Macrophage | Histone;EP300;TF-Binding;POL2;GRO-seq | 114 |
| Mammary_gland | TF-Binding;DHS;Histone;POL2;CAGE | 111 |
| MC3T3-E1 | DHS;TF-Binding;Histone | 31 |
| MEF_E13.5 | TF-Binding;Histone;POL2;MNase;GRO-seq | 47 |
| Megakaryocyte | TF-Binding;Histone;EP300;CAGE | 15 |
| MEL | TF-Binding;DHS;EP300;POL2;Histone | 77 |
| Mesodermal_cell | Histone;TF-Binding;DHS;POL2 | 34 |
| Microglia | DHS;Histone;TF-Binding;CAGE | 17 |
| Midbrain_E11 | TF-Binding;Histone;EP300 | 7 |
| MoDC_BM | DHS;Histone;TF-Binding | 8 |
| MPP | Histone;DHS;TF-Binding | 9 |
| MSC | TF-Binding;Histone;CAGE | 90 |
| Myeloid_cell | DHS;TF-Binding;Histone;GRO-seq | 15 |
| Neural_tube_embryo | DHS;Histone;TF-Binding;FAIRE | 47 |
| Neuron | FAIRE;Histone;POL2;TF-Binding | 19 |
| Neuron_cortical | Histone;TF-Binding;POL2;DHS;GRO-seq;CAGE | 85 |
| Neuron_cortical_E16 | TF-Binding;Histone;POL2 | 76 |
| Neuron_striatal | Histone;POL2;CAGE | 3 |
| NIH-3T3 | TF-Binding;POL2;DHS;FAIRE;Histone;CAGE | 121 |
| NKC_spleen | DHS;EP300;Histone;TF-Binding | 21 |
| NKT | Histone;DHS;TF-Binding | 5 |
| NPC | DHS;Histone;TF-Binding;MNase;FAIRE;POL2 | 83 |
| NSC | TF-Binding;Histone;FAIRE;POL2;GRO-seq | 41 |
| Nucleus_accumben | TF-Binding;MNase;Histone;POL2 | 67 |
| Olfactory_bulb | POL2;Histone;CAGE | 9 |
| Olfactory_epithelium | TF-Binding;MNase;DHS;Histone | 11 |
| Pancreas | TF-Binding;POL2;CAGE | 28 |
| Pancreas_embryo | TF-Binding;FAIRE;CAGE | 8 |
| Pancreatic_islet | DHS;TF-Binding;Histone | 15 |
| PCT4961 | Histone;TF-Binding;POL2 | 5 |
| PDC | Histone;TF-Binding;EP300 | 9 |
| PDC_BM | DHS;Histone;TF-Binding | 8 |
| Peritoneal_macrophage | TF-Binding;DHS;Histone;GRO-seq | 61 |
| Placenta | Histone;POL2;CAGE | 12 |
| Pre-B | Histone;TF-Binding;DHS | 14 |
| Pre-pro-B | TF-Binding;Histone;GRO-seq | 14 |
| Pro-B_BM | Histone;TF-Binding;DHS | 26 |
| Prostate | TF-Binding;Histone;CAGE | 39 |
| Retina_neonate | TF-Binding;Histone;DHS | 45 |
| Rib_chondrocyte_P1 | TF-Binding;Histone;EP300;POL2 | 7 |
| Sperm | DHS;MNase;TF-Binding;Histone | 41 |
| Spermatid | Histone;TF-Binding;POL2 | 14 |
| Spleen | TF-Binding;DHS;Histone;POL2;CAGE | 30 |
| Spleen_leukemia | Histone;POL2;TF-Binding | 7 |
| Stomach_neonate | DHS;Histone;CAGE | 8 |
| Striatum | DHS;Histone;POL2;CAGE | 26 |
| Testis | TF-Binding;POL2;Histone;MNase;CAGE | 122 |
| Th1 | TF-Binding;EP300;Histone;DHS | 59 |
| Th2 | TF-Binding;EP300;Histone | 39 |
| Thymocyte | TF-Binding;Histone;MNase;POL2 | 76 |
| Thymus | Histone;TF-Binding;DHS;POL2;CAGE | 34 |
| Treg_cell | DHS;Histone;TF-Binding | 34 |
| TSC | TF-Binding;DHS;Histone;POL2;FAIRE | 41 |
| Uterus | TF-Binding;POL2;CAGE | 23 |
| V6.5 | TF-Binding;DHS;POL2;Histone;EP300;GRO-seq | 336 |
| WAT | TF-Binding;Histone;POL2;GRO-seq | 32 |
| ZHBTc4 | TF-Binding;DHS;Histone;POL2 | 36 |
